# Partially characterized topology guides reliable anchor-free scRNA-integration

**DOI:** 10.1101/2024.10.22.619682

**Authors:** Chuan He, Paraskevas Filippidis, Steven Kleinstein, Leying Guan

## Abstract

Single-cell RNA sequencing (scRNA-seq) is an important technique for obtaining biological insights at cellular resolution, with scRNA-seq batch integration a key step before downstream statistical analysis. Despite the plethora of methods proposed, achieving reliable batch correction while preserving the heterogeneity of biological signals that define cell type continues to pose a challenge, with existing methods’ performance varying significantly across different scenarios and datasets. To address this, we propose scCRAFT, an autoencoder model designed to segregate cell-type-related biological signals from batch effects for reliable multi-batch scRNA-seq integration. scCRAFT comprises three key loss components: a reconstruction loss that targets observation reconstruction, a multi-domain adaptation loss aimed at eliminating batch effects, and an innovative dual-resolution triplet loss for preserving topology within each batch, which is introduced as an effective mechanism to counteract the over-correction effect of domain adaptation loss amid heterogeneous cell distributions across batches. We show that scCRAFT effectively manages unbalanced batches, rare cell types, and batch-specific cell phenotypes in simulations, and surpasses state-of-the-art methods in a diverse set of real datasets.

## 1 Introduction

The rapid expansion of single cell RNA sequencing (scRNA-seq) has revolutionized biomedical research in the last decade, and is currently reshaping our understanding of cellular diversity and function across multiple biological landscapes. While individual experiments continuously generate new scRNA-seq data, meta-analyses and large-scale collaborations, such as the Human Cell Atlas [1] and Human BioMolecular Atlas Program [2, 3], dedicate substantial effort to effectively integrating data from various settings and conditions, each with distinct technical and biological characteristics. Robust scRNA-seq data integration is challenging due to the interwinding of cellular state heterogeneity and complex batch effects, which may arise from differences in experimental and procedural factors [4, 5], such as sequencing depth and technologies, reagent variations, tissue origin, and inter-individual variability [6–8].

To overcome these integration challenges, a plethora of computational approaches have been developed aiming to maintain a balance between batch mixing and conservation of biological variation [9, 10]. We roughly categorize them into two categories. The first category is anchor-based integration approaches that align anchor cell pairs considered as sharing the same cell type from different batches, with the anchor pairs identified commonly based on mutual nearest neighbors (MNN) [6] or other related techniques such one-way matching [11, 12] and optimal transport [13–15]. Several popular integration methods belong to this category, including BBKNN [16], Scanorama [17] and the widely used Seurat [10]. However, the reliability of anchor-based integration methods rely strongly on high-quality anchor pairs, which can be hampered by limited sharing in cell types across batches [18]. The second category is mixing-loss-based integration approaches, which pair reconstruction loss together with a batch-mixing loss to encourage well-mixed batches if possible. For example, the widely used Harmony [19] and scVI [20] both belong to this category. Harmony is an iterative integration method that minimizes the clustering loss in the embedding space together with batch mixing loss [19], while scVI considers a variational autoencoder (VAE) deep-learning framework and aims to construct batch-free latent embedding with the help from the common prior [20]. The flexibility of deep-learning models and scVI’s framework have been leveraged and led to many variants [21–26]. Despite these exciting progresses, it has been noted that weakly-regularized mixing-loss-based methods, such as scVI, tend to under-correct batch effects whereas strongly regularized mixing-loss-based methods, such as Harmony and scGAN [27], tend to over-correct and remove meaningful signals for defining cell types. As an attempt to alleviate this issue, several deep-learning variants have been proposed to leverage anchor pairs [28–32]. For example, iMAP incorporates a refinement phase utilizing reweighted MNN and generative adversarial networks (GANs) [28]; scDML introduces high-resolution clustering and MNN connectivity between cell clusters to facilitate cluster merging, and finalizes batch correction using deep metric learning. Although these anchor-based refinements achieve more stable embeddings with improved batch-mixing, they inherit the drawbacks associated with MNN techniques.

To address these limitations of existing work, we developed scCRAFT (**sc**RNA-seq batch **C**orrection and **R**eliable **A**nchor-**F**ree integration with partial **T**opology), an anchor-independent deep learning framework with stable training design. scCRAFT employs a synergistic approach that combines the strengths of VAEs, discriminative learning, and a within-batch topology-preserving triplet loss based on dual-resolution clustering. This innovative design prioritizes conservation of desired cellular heterogeneity amid batch-effect removal and offers reliable integration. To assess the efficiency of this approach, we performed large-scale benchmarking analysis in simulated and real-world datasets, featuring various sequencing technologies, imbalanced cell distribution, nested batch effects, and multi-species integration tasks. Our bench-marking results demonstrate that scCRAFT not only consistently outperformed current state-of-the-art methods across all tasks, but also provided good scalability on large-scale datasets.

## 2 Results

### 2.1 Overview of scCRAFT

The scCRAFT framework integrates scRNA-seq data through three key components. First, a custom variational autoencoder (VAE) reconstructs the observed count matrix from a latent embedding Ƶ, ensuring fidelity to the original data. Then, a multi-domain GAN loss is employed for domain adaptation and reducing batch discrepancies within the embedding Ƶ. The final component and a major contribution of scCRAFT is the dual-resolution triplet loss, which encourages the embedding space Ƶ to reflect the within-batch topology of the original dataset 𝒳 (Fig. 1a).

**Fig. 1.**
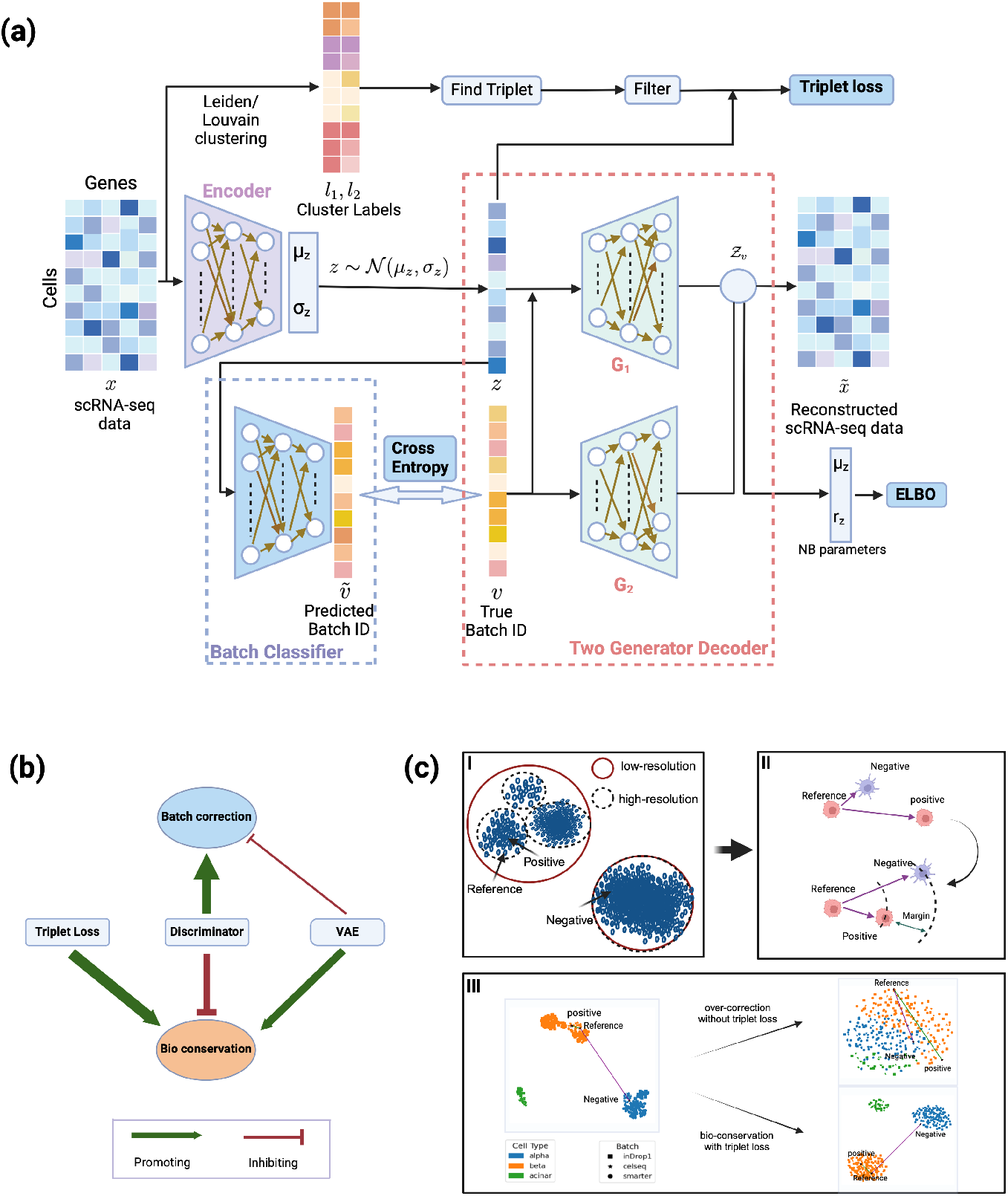
Overview of the scCRAFT pipeline. (a) Schematic diagram of scCRAFT’s architecture. The algorithm includes a variational autoencoder at its core, which integrates a batch-ignorant encoder and dual generator-decoders, trained through an Evidence Lower Bound (ELBO) loss. A batch classifier undergoes adversarial training alongside the encoder utilizing cross-entropy loss. Additionally, the model incorporates triplet loss, calculated within the embedding space Z, guided by labels derived from unsupervised clustering performed on X. (b) Simplified diagram of the conceptual blueprint of scCRAFT, illustrating the synergistic contribution of the VAE, batch classifier, and triplet loss towards the overarching goals of biological conservation and batch correction. (c) I and II illustrate how the triplets are chosen based on the low and high-resolution clustering and the mechanism of triplet loss training respectively. IIII demonstrates the application of scCRAFT on a selected subset from the pancreatic islet dataset, which includes three batches and three cell types. The results compare modeling outcomes with and without the implementation of the triplet loss, and show the positions of an example triplet before and after training.

While observation reconstruction via VAE can produce Ƶ that captures meaningful variations in 𝒳, the basic VAE is often susceptible to batch effects, leading to embeddings that reflect these unwanted variations. The GAN component addresses this issue by training a batch discriminator and encouraging the mixing of embeddings across batches, but can also lead to over-correction in the presence of heterogeneous cellular compositions and blurred distinctions between cell types, especially rare ones. To counter this, we design the dual-resolution triplet loss, which preserves biological signals by maintaining partially characterized yet reliable clustering information within each batch that is independent of any batch effects (Fig. 1b). scCRAFT integrates these components to generate batch-invariant embeddings that also retain the critical biological signals necessary for accurate cell type identification.

The dual-resolution triplet loss is constructed by first performing low-resolution clustering on 𝒳, which organizes cells into clusters of relatively larger sizes. Cells from the same batch but different low-resolution clusters are unlikely to belong to the same cell type. Then, high-resolution clustering is conducted to obtain refined groupings, with cells in the same high-resolution cluster likely sharing the same cell type. Triplet selection within batches is guided by these cluster memberships: each triplet includes a reference cell, a positive cell from the same high-resolution cluster, and a negative cell from a low-resolution cluster. In other words, scCRAFT does not assume exact clustering accuracy but leverages the reliable topology within batches to ensure that cells from highly distinct biological conditions remain separate while preventing the close positioning of cells with very small differences in the embedding space. The combined multi-GAN loss and dual-resolution triplet loss mechanisms encourages the VAE to generate embeddings that effectively remove batch effects while preserving desired biological signals (Fig. 1c, Methods).

### 2.2 scCRAFT demonstrates robustly improved performance across various simulation scenarios and real datasets

We evaluated scCRAFT alongside seven other state-of-the-art integration methods: scVI, Harmony, Seurat, Scanorama, BBKNN, iMAP, and scDML (Suppl section 2). All methods were assessed based on their integration accuracy and scalability in terms of computation efficiency and peak memory usage. Integration accuracy was assessed using a suite of metrics divided in two categories. The first category consisted of four metrics which are averaged to be the bio-conservation score, measuring each method’s ability to preserve biological variance by examining the accuracy of global clustering and the distances between clusters. The second category consisted of six metrics which are averaged to be the batch-mixing score, measuring the uniformity of integration across batches to ensure that they are indistinguishable after integration. The composite score was calculated as a weighted average of bio-conservation and batch-mixing, with a 60/40 ratio favoring the former over the latter [8]. For each dataset, we calculated individual scores for each metric, average scores for bio-conservation (NMI, ARI, ASW cell, cLISI) and batch-mixing (ASW batch, PCR batch, Graph Connectivity, kBET, bLISI, True Positive Rate) separately, and composite scores combining these two categories. Finally, an overall score was determined by calculating the average of composite scores from all datasets. We also included one supplementary metrics F1 LISI, which simultaneously evaluated biological conservation and batch mixing and further supported the accuracy of benchmarking (Suppl Section 5). Since BBKNN produces only a two-dimensional embedding, which may bias certain evaluation metrics, we excluded its evaluation scores from the benchmarking. However, we included its UMAP output in Supplementary Figures (Fig. S1-S19), where it showed noticeably lower integration quality compared to scCRAFT and other top performers.

We evaluated each of the methods across nineteen datasets, ten simulated and nine real. These datasets were specifically selected to reflect a broad spectrum of integration challenges (Suppl Section 1). The simulated datasets included scenarios with unbalanced cell counts, batch-specific cell types and rare cell types, and were instrumental in evaluating our method in a controlled setting where the ground truth was known, allowing for a full characterization of batch effects. These datasets encompassed one simple and three complex scenarios: (1) cell types with identical counts evenly distributed across batches; (2) unbalanced batch sizes, where the size of batch 1 was reduced to represent 50% (batch1-0.5), 25% (batch1-0.25), and 12.5% (batch1-0.125) of its initial size; (3) batch-specific cell types, where the overlap of cell types across batches was modified to include only 5 (common-5), 3 (common-3), or 1 (common-1) shared types; and (4) rare cell types, with the proportion of cells in Group1 in each batch reduced to represent 50% (rare1-0.5), 20% (rare1-0.2), and 10% (rare1-0.1) of their initial count. Across all simulated datasets, scCRAFT consistently outperformed all other tested methods in terms of composite scores, followed by Scanorama and, depending on the specific dataset, by scVI or Seurat. In terms of bio-conservation metrics, scCRAFT also achieved the highest average scores and performed comparably or better than all other methods in individual metrics. Specifically, scCRAFT outperformed Seurat by 3.4%, Scanorama by 5.4%, and scVI by 26.6% using the average scores across datasets, and Seurat by 16.8%, Scanorama by 22.9%, and scVI by 25.8% when measured by ASW label, which is the most challenging metric on these simulated experiments (Fig. 2a-b, Fig. S1-S10). On the other hand, while scCRAFT outperformed Scanorama by 9.7%, scVI by 4.5%, and Seurat by 22.9% using the average batch-mixing scores across datasets, the individual batch-mixing score per dataset presented a more complex picture (Fig 2b, Fig S1-S10). scCRAFT achieved the highest average batch-mixing score in eight out of ten datasets, ranking second in the remaining two datasets (“batch1-0.125” and “batch1-0.250”). Further examination revealed that the second-place scores on these two datasets were due to a lower bLISI score in these highly unbalanced batch scenarios (Fig. S2-S3). Despite this, scCRAFT consistently scored highest or comparably in all other batch-mixing metrics, and the embedding plots showed that cell types from different batches were still correctly clustered together. The supplementary F1 LISI metric also supported this conclusion, with scCRAFT ranking first in seven tasks, second in one, and mid-range only for “batch1-0.125” and “batch1-0.250”, where its bLISI score was not top-ranking as previously discussed (Fig. S1–S10). Overall, scCRAFT demonstrated robust improvement in integration quality compared to the baseline methods. Notably, scCRAFT also successfully identified rare cell types in the rare cell type simulation “rare-0.1”, correctly pairing them with corresponding rare cells from other batches (Fig. 2a).

**Fig. 2.**
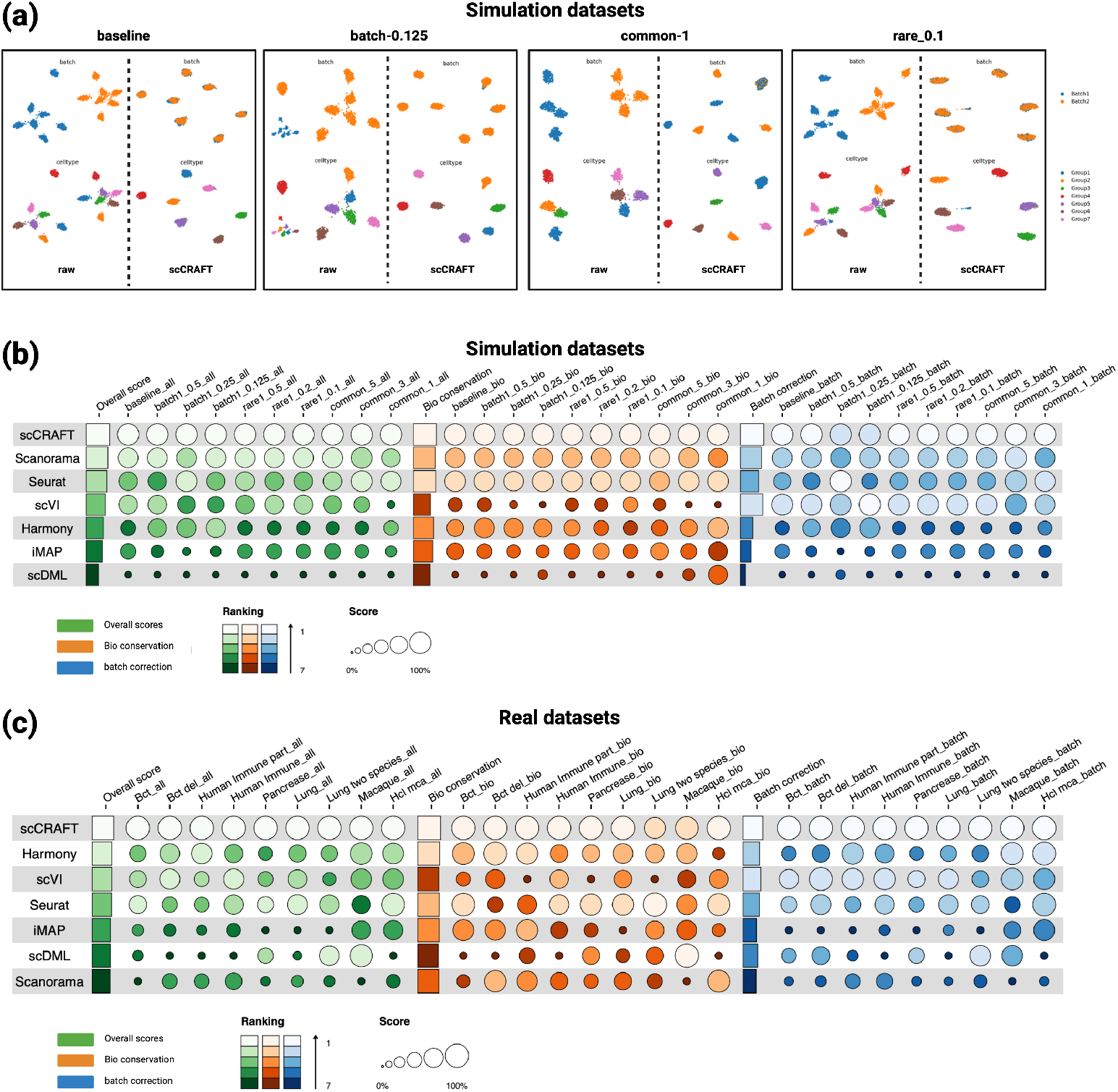
Benchmarking results on simulation and real data. (a) UMAP visualization contrasting unintegrated simulation datasets with scCRAFT-integrated counterparts under various simulation tasks, including baseline where cell types are shared equally across batches of the same size, unbalanced batch sizes (batch-0.125), batch-specific cell types (common-3), and rare cell types (rare-0.1). The upper part of each box shows the integration of different batches and the lower part shows the delineation of cell types (Groups) within these integrated datasets. (b), (c) Overview of the quantitative evaluation of 10 simulations tasks (b) and 9 real datasets (c), including the specific rankings and scores for each evaluation metric across 7 benchmarking methods. The average scores across individual metrics for each dataset are illustrated with orange and blue circles. The average scores across all datasets are illustrated with orange and blue bars. The composite scores for each dataset are illustrated with green circles, and were calculated as the weighted average of bio-conservation and batch mixing, with a 60/40 ratio favoring the former over the latter. The overall score represents the average of the composite scores of all datasets. The diameter of each circle represents a linear rescaling of its corresponding values to a range between 0.1 and 1. The length of each bar represents the absolute average value. Methods are ranked based on their overall score, from highest to lowest.

We subsequently applied scCRAFT to integrate nine real datasets with varied characteristics: small scale datasets with few cell types (bct and bct-del); large, widely used datasets (pancreatic islets, lung atlas, human immune datasets, and a subset of the human immune datasets); a dataset with a large number of batches (macaque dataset); and datasets including data from two-species (lung two species and human mouse datasets) (Suppl Table S1). These datasets presented complex integration scenarios, including nested batch effects arising from varying techniques and donors, integration across different tissues and species, rare cell types and large-scale datasets, accommodating up to 300,000 cells and up to 30 batches. For real datasets, the reference truth was determined by the original annotations provided by the source datasets which have also been adopted in other benchmarking studies [7, 8, 29, 30].

Similar to the results on simulated datasets, scCRAFT consistently achieved the highest composite scores across all real datasets. The performance of other methods varied significantly compared to the simulated datasets, with Harmony, scVI or Seurat emerging as the second-best method depending on the dataset analyzed (Fig. 2c). In terms of bio-conservation metrics, scCRAFT marked the highest average scores in all but two out of nine datasets where it ranked the second best (“Lung two species” and “Macaque”), with Harmony 3.6%, Seurat 5.8%, and scVI 7.5% lower than scCRAFT using the average bio-conservation score across datasets (Fig 2c, Fig S11- S19). In the “Lung two species” dataset, the slightly lower bio-conservation average score was attributed to modestly lower scores in all individual metrics compared to Seurat (1.1% better than scCRAFT), which achieved the best bio-conversation score on this dataset (Fig. S15). In contrast, the individual bio-conservation scores in the “Macaque” datasets were comparable to those of the best-performing method, scDML, whose average bio-conservation score was 1.6% better than scCRAFT (Fig. S16). However, scDML’s performance was considerably lower on several other datasets. As for the batch-mixing metrics, scCRAFT led with the highest average scores on all nine datasets, surpassing the second-best method in this category by an average of 11.5%, with varying degree of improvements compared to the second-best method for different datasets. scCRAFT ranks highest in 8 out of 9 datasets by the supplementary F1 LISI metric, confirming its broad improvements over the baseline methods (Fig. S11–S19).

### 2.3 Successful identification of rare cell types

To evaluate scCRAFT’s effectiveness in identifying rare cell types, we investigate more closely the human pancreatic islet dataset, which is comprised of 16,382 cells which have a significant cell proportion disequilibrium. This dataset is derived from a collection of nine sub-datasets generated through diverse sequencing technologies, including CEL-seq, CEL-seq2, Fluidigm C1, SMART-seq2 and inDrop. The samples cover a broad spectrum of 14 pancreatic cell types, including both pancreatic and immune cells. Despite the balanced batch distribution for each cell type, some cell types such as alpha cells (33.53%), beta cells (25.45%), and ductal cells (13.08%) are highly abundant, while others like macrophages (0.48%), mast cells (0.26%), epsilon cells (0.20%), Schwann cells (0.15%), and T cells (0.04%) are extremely rare (Fig. 3a), making it an excellent case for evaluating how well integration methods can identify and integrate rare cell populations within a mixed-cell population.

**Fig. 3.**
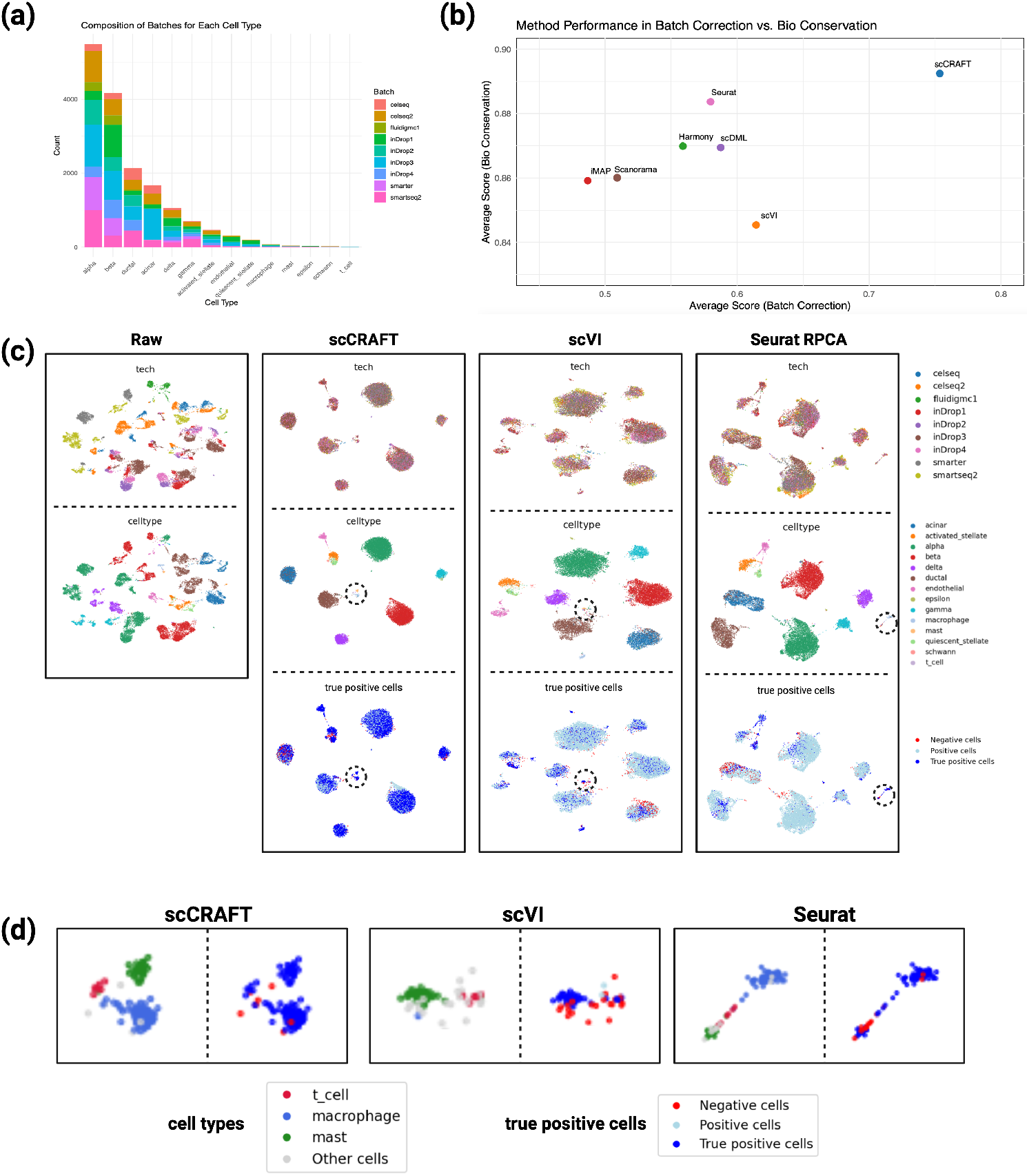
Benchmarking results on pancreatic islet dataset. (a) Bar plot showing the absolute cell count for each cell type and the distribution of batches across different cell types. For visual clarity, only scVI and Seurat are included in the visualization, as scDML was unable to identify rare immune cells. (b) Scatterplot of average bio-conservation score against the average batch correction score for each method. (c) UMAP visualization contrasting the unintegrated pancreatic islet dataset with its integrated counterparts using scCRAFT and two other most popular benchmarking methods. The top panel shows the integration of different batches (sequencing technologies), the middle panel shows the cell types, and the bottom panel shows the distribution of negative, positive, and true positive cells. (d) UMAP visualization focusing on the integration of rare cell types (T cells, macrophages, and mast cells) for scCRAFT, scVI, and Harmony. For each method, the left side of the panel shows the cell type distribution, and the right side shows the classification of cells into negative, positive, and true positive.

scCRAFT effectively integrated pancreatic islet datasets across six different technologies, achieving the highest composite score, followed by Seurat, scDML and scVI. Although the margin of its superiority in individual bio-conservation metrics was relatively modest compared to the second-best method in this category (Seurat), its performance in batch-mixing metrics was consistently higher than other methods (Fig. 2c, Fig. 3b, and Fig. S17b). We then investigated the ability of each method to identify rare cell types, which may not necessarily be accurately reflected on the overall bio-conservation or batch-mixing score. Using UMAP visualizations to assess clustering, scCRAFT was notably effective in identifying and merging very rare immune cell types, including macrophages, mast cells and T cells, while clearly separating them from other pancreatic cells. This task was only partially achieved by other methods (Fig. 3c-3d). We quantitatively evaluated each method’s ability to identify rare cell types by measuring the achieved true positive rates in these types (Fig. 3c-3d). A cell is classified as positive if it is surrounded by cells of the same type and negative if not, and further as a true positive if it is a positive cell with surrounding batch distribution matches the global batch distribution for that cell type [28]. Among the rare cell types previously mentioned, scCRAFT outperforms other methods in identifying neighboring cells of the correct type with only 3.5% negative cells compared to 32.2% for scVI and 16.25% for Seurat, and it also effectively corrects for batch effects among rare cell types, achieving a true positive rate of 91.5%, which is substantially higher than all other methods (Fig. 3c-3d, Fig. S17).

### 2.4 Complex integration across multiple sources of variation

To further assess scCRAFT’s performance in more complex scenarios, we used the lung atlas dataset, originally curated by Vieira et al [33]. This dataset includes 32,472 cells from lung transplant and airway biopsy specimens derived from 16 different donors, sequenced using both 10X Genomics and Drop-seq technologies. Its complexity and strong batch effects arise from several factors, including high interdonor variability, different sequencing protocols without sample multiplexing, and large heterogeneity in cell composition and abundance related to the sampled spatial location, specifically lung parenchyma transplants and upper airway biopsies (Fig. 4a). The combination of multiple sources of technical and biological variation make this dataset a good candidate to assess the robustness and effectiveness of scRNA-seq data integration methods in addressing complex integration challenges.

**Fig. 4.**
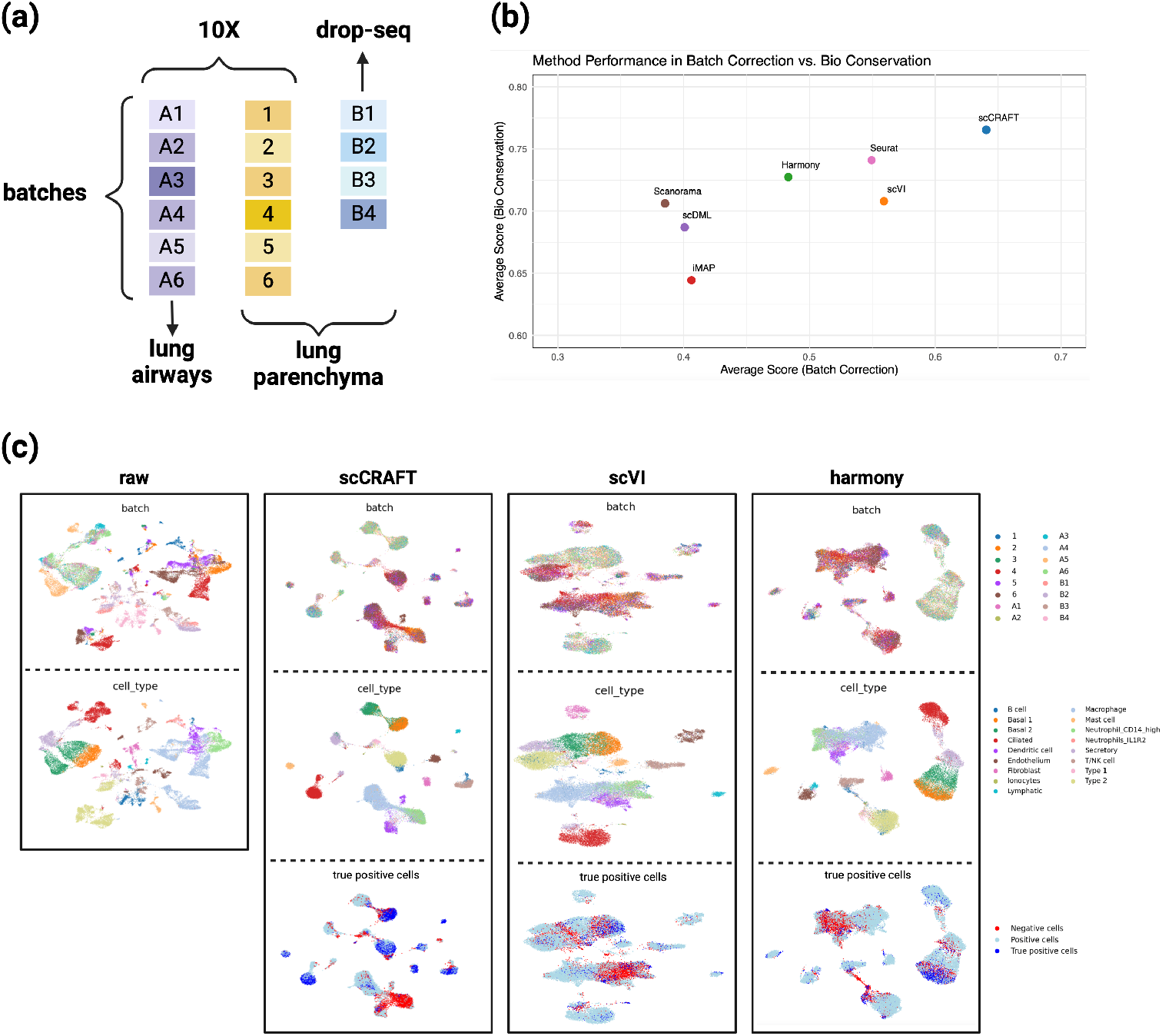
Benchmarking results on lung atlas dataset. (a) Batch composition and nested batch effects in the lung atlas dataset, including three studies, two sequencing technologies, and two tissue types. (b) Scatterplot of average bio-conservation score against the average batch correction score for each method. (c) UMAP visualization contrasting the unintegrated pancreatic islet dataset with its integrated counterparts using scCRAFT and two other most popular benchmarking methods. The top panel shows the integration of different batches (samples), the middle panel shows the cell types, and the bottom panel shows the distribution of negative, positive, and true positive cells.

scCRAFT clustered the cells into groups compatible with the cell types and substantially reduced technical batch effects, outperforming other tested methods in terms of composite and average bio-conservation and batch-mixing scores (Fig. 4b, Fig. S18b). Notably, scCRAFT differentiated “Type 2” cells from B cells, particularly those originating from donor B4, and distinguished “Neutrophil CD14 high”, macrophages, and dendritic cells from one another, tasks that most other integration methods struggled with (Fig. 4c, Fig. S18a). Notably, scCRAFT outperformed the three top-performing baseline methods—Seurat, scVI, and Harmony—by a large margin in terms of true positive rate, achieving a rate of 0.27, which is considerably higher than Harmony (0.09), scVI (0.08), and Seurat (0.1) (Fig. 4b-4c). These results highlight scCRAFT’s effectiveness in handling challenging integration tasks in the presence of significant batch effects.

### 2.5 Cross-species integration with low cell type overlap

We examined scCRAFT’s integration capabilities on a large dataset comprising 372,789 cells and two batches, derived from the Human Cell Landscape (HCL) [34] and Mouse Cell Atlas (MCA) [35]. This dataset poses a unique challenge due to the low overlap in cell types between the two species. Specifically, the HCL batch includes 249,845 cells spanning 63 cell types, and the MCA batch includes 122,944 cells spanning 50 cell types, with only 18 cell types shared between the two. These characteristics make this dataset suitable for testing scRNA-seq cross-species data integration, with potential applications extending to comprehensive atlas-level studies.

Overall, scCRAFT ranked first in both average and most individual bioconservation and batch-mixing scores, with Seurat following closely behind (Fig. 5a-c). Of note, the absolute average bio-conservation scores across all methods were consistently lower for this dataset (range 0.6 - 0.67), compared to the pancreatic islet dataset (range 0.84 - 0.89) and the lung atlas dataset (range 0.65 - 0.77) (Fig. 3d, 4d, 5b). Cell type delineation was challenging for all methods. Still, compared to scCRAFT, scVI, Scanorama and scDML showed worse performance for batch-mixing and exhibited a stronger tendency to separate identical cell types into multiple clusters, potentially complicating subsequent cell type annotation. Conversely, Harmony, BBKNN, and iMAP tended to merge different cell types compared to scCRAFT due to over-correction, obscuring true biological signals for cell type identifications (Fig. 5a-c, Fig. S19). These findings support the potential of scCRAFT to integrate datasets from different species with low numbers of shared cell types.

**Fig. 5.**
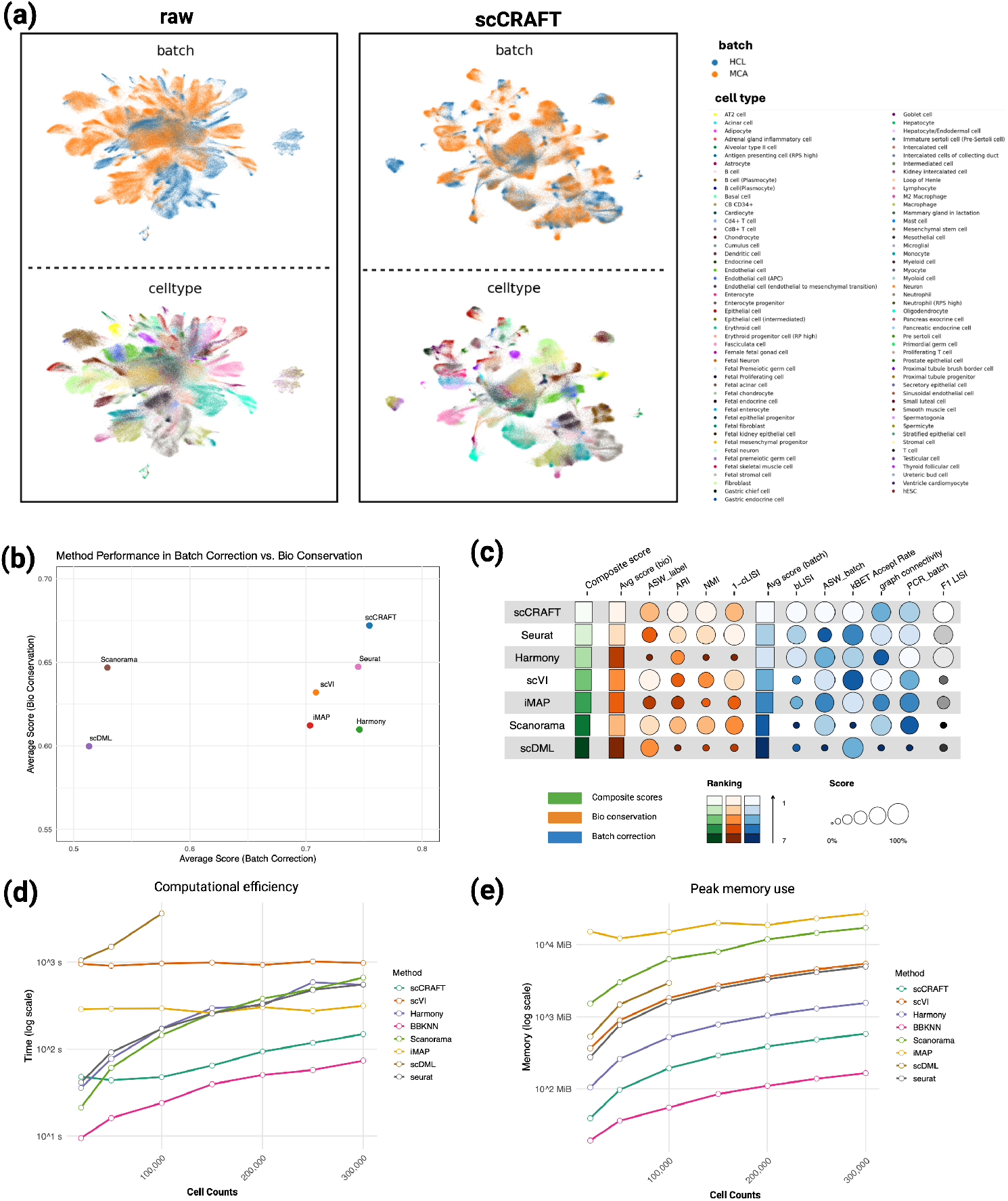
Benchmarking results on human and mouse dataset. (a) UMAP visualization, contrasting the unintegrated dataset with its scCRAFT-integrated counterpart. (b) Scatterplot of average bioconservation score against the average batch correction score for each method. (c) Overview of the quantitative evaluation, including the specific values and rankings for each evaluation metric across the different methods. The individual scores for each metric are illustrated with orange and blue circles. The average scores across all metrics are illustrated with orange and blue bars. The composite score for each method is illustrated with a green bar, and was calculated as the weighted average of bio-conservation and batch mixing, with a 60/40 ratio favoring the former over the latter. Due to the high computational expense associated with calculating postive/true positive rates, we exclude them for this experiment. (d) Runtime with increasing cell counts for each integration method in a subset of the human-mouse dataset. (e) Peak memory usage with increasing cell counts for each integration method in a subset of the human-mouse dataset. Note: scDML does not have evaluations for large numbers of cells because the computation time becomes prohibitively high.

### 2.6 Scalability and computational efficiency in large data

We evaluated scCRAFT computational performance by measuring total runtime and peak memory usage, using an Intel 64-kernel CPU. To this end, we downsampled the combined Human Cell Landscape and Mouse Cell Atlas dataset into seven benchmark datasets of increasing sizes (from 20,000 to 300,000 cells). scCRAFT exhibited a modest rise in total runtime beyond 100,000 cells, yet it remained the second-best method, only surpassed by BBKNN, which performed significantly worse in the embedding. In contrast, the runtime of other deep learning-based methods, like scVI and iMAP appeared largely independent of cell numbers due to the early stop (scVI) or batch-wise sampled training (iMAP), with scVI and iMAP exhibiting, on average, ten-fold and three-fold longer runtimes than scCRAFT, respectively. Harmony, Scanorama, and Seurat also showed a nearly linear increase in runtime with increasing cell counts, slightly outperforming scCRAFT at scales below the 50,000, though the differences were minimal (48s for scCRAFT, 41s for Seurat, 36s for Harmony, and 21s for Scanorama). In terms of peak memory usage, most methods displayed a linear increase with growing cell numbers. At 300,000 cells, BBKNN had the lowest peak memory usage (around 150 MiB), followed by scCRAFT (500 MiB) and Harmony (1,500 MiB). iMAP and Scanorama had the highest peak memory usage (over 10,000 MiB), followed by scVI and Seurat (around 5,000 MiB) Additional discussions on the time and memory story complexity can be found in Supp Section 4. These results demonstrate scCRAFT’s computational efficiency, making it suitable for large-scale scRNA-seq data integration tasks.

## 3 Discussion

scRNA-seq technologies and experimental designs are rapidly evolving, highlighting the need for flexible integration methods capable of handling multiple sources of technical variation. However, correcting strong batch effects comes with the risk of obscuring relevant biological signals, which underlines the importance of optimizing the trade-off between bio-conservation and batch-mixing. scCRAFT provides a computationally efficient and innovative framework for more reliable scRNA-seq data integration, designed to overcome the limitations of anchor-based MNN methods and conventional VAE modeling. Its architecture uniquely combines a VAE, enhanced by a multi-domain conditional GAN, with a dual-resolution triplet loss to conduct reliable scRNA-seq integration under partially characterized batch-free topology. This configuration effectively captures biological signals and generates batch-invariant embeddings, thereby achieving improved balance between batch correction and conservation of biological variation. While these three core components–VAE, GAN, and dual-resolution triplet loss–are indispensable for its functionality, scCRAFT remains robust even as the loss proportions of these components vary widely, as demonstrated through ablation studies and tuning experiments (Section 4.3, Fig. S20-21).

The dual-resolution triplet loss mechanism in conjunction with the VAE training approach represents a substantial advancement in scRNA-seq data integration techniques. Traditional triplet loss-based methods typically adopt the triplet loss to remove the batch effects, by utilizing the embeddings obtained from the VAE or other steps to identify reference and positive cells across batches [29, 31, 32] and refine the embeddings by drawing similar cells from different batches to be closer to mitigate batch effects. A key limitation of this traditional strategy is that the triplets are derived from initial embeddings, which may not always be reliable to identify similar cells as identifying similar cells across batches is a most challenging task in scRNA-seq integration. In contrast, scCRAFT’s strategy only leverages the triplet loss to capture the topology of raw counts from individual batches trained along with the VAE for better biological conservation in the embedding. This approach circumvents the need to identify triplets across batches based on potentially unreliable preliminary embeddings. Moreover, the dual-resolution mechanism allows the distinction between confident grouping and unsure assignment, and adeptly preserves the hierarchical structures inherent in biological data, which is crucial for high-resolution clustering analysis after integration. This dual-level resolution is key for detailed cell type annotation and enhances the ability to identify and analyze cell subtypes within larger clusters. In the framework of scCRAFT, this dual-resolution triplet loss effectively mediate the over-correction tendency of the multi-domain GAN component that undertakes the batch-removal task, which has been shown as an efficient approach for domain alignment, but can also erroneously align samples in the presence of heterogeneous signal across domains.

In our comprehensive benchmarking analysis, we evaluated scCRAFT alongside six other widely-used integration methods. Performance of the six existing methods was data dependent and varied significantly across different test datasets. For instance, in simulated scenarios, Seurat demonstrated strength in biological conservation and Scanorama showed a slight edge in batch mixing, whereas Harmony struggled with both tasks, particularly in the presence of batch-specific clusters and rare cells. In real datasets, scVI, Harmony, and Seurat were markedly better in batch-mixing compared to iMAP, scDML, and Scanorama, although their performance in bio-conservation was less consistent.

One limitation of the benchmarking process is the reliance on source-provided cell annotations as the ground truth for evaluating biological conservation. These annotations may not be entirely reliable, and annotation standards can vary across datasets. Despite these inconsistencies, the datasets used are widely adopted and individually validated in each benchmarking study, making them the most dependable available. Another limitation is the hyperparameter tuning for different loss components. While we have shown that default hyperparameters can generally be used across datasets and provided a heuristic method for tuning without true cell type annotations, like other scRNA-seq integration methods with hyper-parameters, a more automatic and adaptive approach to balance different loss components could be beneficial.

Despite the limiations, scCRAFT demonstrates consistently robust performance across various technical and biological contexts, significantly improving batch-mixing and bio-conservation through its novel dual triplet loss approach. Its scalability and computational efficiency make it particularly suitable for large, complex scRNA-seq datasets and atlas-level single-cell studies, even with limited computational resources. Looking ahead, the inherent flexibility of scCRAFT’s deep learning architecture makes it well-suited for an array of additional tasks including semi-supervised integration. Notably, scCRAFT’s anchor-free advantage also offers opportunities for a more general framework that could be extended to multi-omic datasets [36, 37]. These promising avenues could significantly enhance scCRAFT’s utility in the ever-expanding landscape of single-cell and multi-omic data integration.

## 4 Method

### 4.1 The scCRAFT framework

The scCRAFT aims to construct batch-invariant embedding that well-capture the heterogeneous cell type compositions from different batches using a tailored variational autoencoder (VAE) architecture that consists of three loss components: 1) the reconstruction loss, 2) the domain adaptation loss based on a multi-domain GAN, and 3) the dual-resolution triplet loss that counteracts the mult-domain GAN’s over- correction effect. Here, we describe scCRAFT with implementation details deferred to Suppl Section 3.

#### 4.1.1 Variational autoencoder and the reconstruction loss

The encoder, *E*, is a multi-layered feed-forward neural network which consists first of a network of size (*p* → 1024 → 512) where *p* is the dimension for the input expression profile *X*. This network is then followed by two separate output layers of size *p*_0_ to yield the posterior mean *q*_*η*_ and posterior variance *q*_*ϵ*_ of latent embedding *Z* which is assumed to be generated from a standard normal prior (*p*_0_ = 256 by default) and whose posterior distribution is also modeled as normal: Ƶ ∼ *N* (*q*_*η*_, *q*_*ϵ*_). Batch normalization and ReLU activation functions are employed following each dense layer to stabilize the learning process and introduce non-linearity.

The decoder network *G* comprises two steps. The first step incorporates the latent variables Ƶ and batch information vectors *v* to construct a batch-integrated representation of *p*-dimensions. It utilizes a composite structure where the first decoder *G*_1_ decodes the latent space Ƶ concatenated with the batch effect vector *v*, followed by a the network *G*_2_ decoding only the batch effect vector. These two outputs are then combined and activated through a ReLU function to generate the final batch-integrated representations 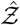 of *p*-dimensions:

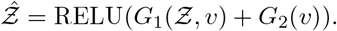

This construction encapsulates the complex interplay between biological variation and technical noise. Then, assuming a negative binomial (NB) distribution for the input observations, the second step determines the mean and dispersion parameters 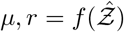, where *f* (·) is a fully connected neural network together with an exponential transformation (keep positive) that connects the latent representation 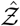 to the parameters *μ* and *r*. For all *x* ∈ 𝒳 denoting the gene expression profile and (*μ, r*) be its corresponding distribution parameters, we have

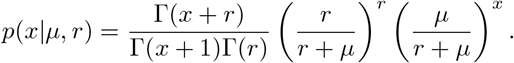

The training of the traditional VAE is guided by the Evidence Lower Bound (ELBO), which is defined below

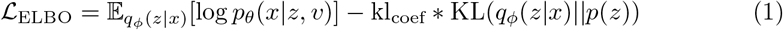

where *ϕ* and *θ* represent all parameters of the encoder network and decoder network, respectively. *q*_*ϕ*_(*z*|*x*) is the encoder’s posterior distribution, *p*(*z*) is the prior distribution of *z* and *p*_*θ*_(*x*|*z, v*) is the likelihood of the observed data given the latent variables, parameterized by the negative binomial distribution for capturing the count data’s overdispersion. The KL divergence term encourages the posterior distribution to align with the prior, whereas the expected log-likelihood term aims to maximize the data’s reconstruction accuracy. Note that kl_coef_ = 1 in the standard variational inference framework. However, this can lead to severe over-correction, and kl_coef_ is often set much smaller than 1 for scRNA-seq integration, e.g., kl_coef_ = current epoch*/*400 in scVI with the default number for training epochs smaller than 400. Since we also have the multi-domain GAN for batch correction, we choose a smaller KL divergence coefficient kl_coef_ = 0.005 for all datasets, which is fixed throughout all our experiments.

The NB loss may lead the model to overly focus on individual gene expression. Hence, we also incorporate the cosine similarity as another reconstruction loss to help better preserve the correlation across genes for two cells:

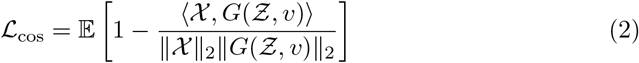

where *v* is the one-hot encoded batch vector, and ⟨·, ·⟩ denotes the dot product between the original and the reconstructed gene expression profiles.

#### 4.1.2 Domain adaptation via multi-domain GAN

To enhance the ability to conduct batch-correction for multiple batches, we include a multi-domain GAN component to encourage the alignment of different batches. The multi-domain GAN consists of a discriminator separating different batches as a batch classifier and the associated loss that encourages our embedding to fool this classifier. The discriminator network, denoted as *D*, updates its parameters by minimizing the cross-entropy loss, to find the most discriminative direction for assigning the current Ƶ into their corresponding batches. Concurrently, the encoder is trained to maximize this loss, thereby encouraging the generation of batch-invariant embeddings Ƶ. In other words, define the cross-entropy loss function for |*S*| batches as below:

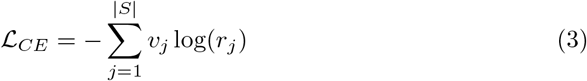

Here, *v*_*j*_ is the true label for the batch, and *r*_*j*_ is the predicted probability (softmax output) of from *j*th batch. The batch-classifier (discriminator) aims to minimize this loss while the encoder seeks to maximize the cross-entropy loss so that the classifier cannot easily differentiate between batches. This adversarial training forms a minimax game defined by the objective:

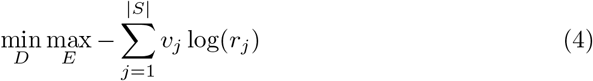

This model component encourages batch-ignorant embeddings such that scCRAFT is not constructing Ƶ driven by batch effects.

#### 4.1.3 Dual-resolution triplet Loss

To effectively address the challenge of conservation of biological variation during batch correction, scCRAFT introduces a dual-resolution triplet strategy. The triplet loss aims to encourage that embeddings of samples from the same biological cluster are positioned closer together, while those from different clusters are distanced. Specifically, the triplet loss is defined as:

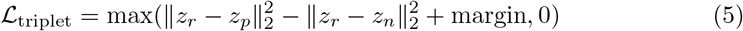

where *z*_*r*_, *z*_*p*_, and *z*_*n*_ represent the embeddings of the reference, positive, and negative cells, respectively, with positive cells defined as those believed to be in the same cell type as the reference cell and negative cells defined as those believed to be from different cell types. This strategy is designed to enhance the model’s ability to accurately capture and represent the intricate biological diversity present within single-cell datasets. The margin was set to be 5 across experiments.

A major challenge in the triplet loss is identifying positive and negative cells given a reference cell. A widely accepted rule is that cells from the same cluster in a given batch tend to be from the same cell type. However, defining cell clusters is subjective: in the standard pipeline, cell clusters are identified using the Leiden algorithm given a user-specified resolution parameter. When the resolution parameter is large, we identify many small clusters with the risk of over-partitioning. When the resolution parameter is small, few clusters are identified with the risk of merging the same cell type into one big cluster. In practice, it requires careful thought to determine a good resolution parameter. Further, it is often the case that a fixed resolution cannot recover all cell types of interest. To resolve this challenge, we introduce a novel dual-resolution triplet loss that operates as follows:

- **Lower Resolution Clustering:** Initially, cells are clustered into broad categories using the Louvain algorithm [38] or Leiden algorithm [39] with the default resolution of 0.5, which are both unsupervised labeling techniques. This step aims to identify large and distinct biological clusters, setting the stage to find the initial triplets. We identify the reference, positive (same cluster with reference), and negative cells (different cluster from reference) based on this low-resolution clustering (Fig. 1c).
- **Higher Resolution Clustering:** Subsequently, within each broad cluster established in the lower resolution step, a higher resolution clustering (default resolution 7) is executed to uncover and delineate the subtypes present. At this stage, the initial triplets are further filtered to include only those with reference-positive pairs confirmed to belong to the same subtype (same high-resolution cluster). This precision ensures the model does not apply the triplet loss to pairs from different high-resolution groups, thereby maintaining the integrity and distinction of subtle biological variations.

In the execution of our dual-resolution strategy, it is important to note that the triplet loss calculation is constrained to within individual batches, hence, not influenced by the batch effects.

#### 4.1.4 Optimization of the scCRAFT Framework

In our scCRAFT framework, the training process is governed by a carefully designed optimization algorithm that balances several objectives, aiming to enhance the biological relevance of the learned representations while mitigating batch effects. The optimization objectives for the encoder and generator, as well as the discriminator, are detailed below.

The encoder *E* and generator *G* are jointly optimized to minimize a composite loss function that integrates various regularization terms:

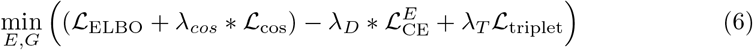

where 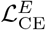 represents the adversarial loss against the discriminator, and is defined as:

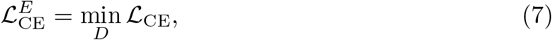

with ℒ_CE_ being the cross-entropy loss separating different batches and *D* being the discriminator aiming for the best separation. ℒ_ELBO_ is the combined VAE loss, ℒ_triplet_ is the triplet loss function that enhances separation between different cell types, and ℒ_cos_ is the cosine similarity loss.

We adopt the learning rate 0.001 for both updating the encoder *E*, generator *G*, and the discriminator *D* (in ℒ_CE_), with the discriminator undergoing ten updates for every one update of *E* and *G*. To ensure uniform batch representation, we composed each training mini-batch of 512 cells by mixing an approximately equal number of cells from each batch. Additionally, we implemented a default 50 warm-up epochs where cross-entropy loss 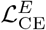 was excluded, allowing the embedding Ƶto capture inherent data patterns prior to integrating the cross-entropy loss, which facilitates batch mixing.

### 4.2 Hyperparameter Setting

In our composite loss function, the proportions of various loss components are kept constant across both simulations and real datasets. Specifically, we assign the coefficients for the discriminator loss, triplet loss, and cosine similarity loss as *λ*_*D*_ = 0.2, *λ*_*T*_ = 1, and *λ*_cos_ = 20, respectively. We are particularly interested in learning the contribution and robustness of the two components of our scCRAFT framework that actively remove batch effects and maintain batch-free topology - the multi-domain GAN and triplet loss.

To evaluate the contribution of these two components, we first conducted an ablation study by setting the parameters *λ*_*D*_ and *λ*_*T*_ to zero in the pancreas and lung integration tasks. Subsequently, we quantitatively evaluated the impact of these components on integration performance by incrementally adjusting *λ*_*D*_ from 0 to 0.7 and *λ*_*T*_ from 0 to 5 to demonstrate the model’s robustness in a wide range of parameter values (Fig. S20a-b).

While the parameter settings adopted in our experiments generally demonstrate robust performance, we also acknowledge the potential for suboptimal outcomes in specific scenarios where users might require tailored hyperparameter tuning. In realworld applications, the absence of cell type annotation often complicates the evaluation of integration performance and hyperparameter tuning. To address this challenge, we have developed a fast strategy to quantify the evaluation suggestion raised by a recent benchmarking paper for assessing biological conservation [40]. Specifically, we utilize low-resolution clusters within each batch as pseudo true labels for post-integration evaluation. Post-integration, we calculate NMI and ARI for each batch using these labels, and then average them across all batches. Intuitively, these metrics serve as proxies of whether major biological variations within individual batches are preserved following integration. Notably, the trends observed in the averaged NMI and ARI closely parallel the variations in the “real” metrics when tuning the parameters of *λ*_*D*_ and *λ*_*T*_, calculated using known cell type labels on the integrated dataset (Fig. S20b- c). This close correlation between the unsupervised and true metrics highlights the effectiveness of our approach in evaluating integration quality, offering a dependable method for tuning scCRAFT’s parameters and selecting integration methods without relying on annotations.

The clustering resolution for triplet identification is set at a default high value of 7. Notably, the low-resolution parameter, pivotal for primary cluster formation, necessitates careful tuning. A value of 0.5 is empirically robust for most datasets; however, for simulations or datasets with more detailed sub cell types, like the lung atlas dataset and HCL-MCA dataset discussed, a higher resolution of 1 is adopted to delineate them and may be advantageous. Conversely, for simpler annotations like the Bct and Bct-del datasets, a lower resolution of 0.2 ensures the merging of the same cell types. The integration results for the mentioned datasets with alternative resolutions are shown in Fig. S21.

Regarding clustering algorithms, Leiden and Louvain perform comparably. Leiden may offer superior results in certain cases but is slower, especially with large datasets. Thus, while Louvain is recommended for efficiency, Leiden is worth considering when optimal results justify the computational expense.

### 4.3 Data Preprocessing

Our scRNA-seq data preprocessing follows the established Scanpy pipeline [41]. Postimport of raw count matrices into Scanpy AnnData objects, low-quality cells with fewer than 300 detected genes and genes found in less than 5 cells were filtered out to prevent the misleading alignment of cells characterized by significant dropout events with those exhibiting low transcriptional activity [17]. Subsequently, each cell’s library size was normalized to 10,000 reads, a process involving the scaling of gene counts by total counts per cell and adjustment by a factor of 10,000. This step was followed by a log transformation using log(1 + *x*), ensuring a stabilized variance across the data. Additionally, we identified the top 2000 highly variable genes, crucial for capturing the dataset’s biological diversity while accounting for batch effects. This preprocessing was applied uniformly across all benchmark methods.

## Acknowledgements

This work is supported by NSF DMS2310836 and 5U01AI167892.

## Declarations

- Data availability: See Supplementary Material, Suppl Section 1
- Code availability: https://github.com/ch2343/scCRAFT/tree/main

