## Supplementary file for "Partially characterized topology guides reliable anchor-free scRNA-integration"

### Contents

**1 Dataset details** **3**
1.1 Simulation datasets . . . 3
1.2 Real dataset . . . 4

**2 Benchmarking Methods** **8**

**3 Detailed algorithm** **9**

**4 Additional discussion on computational cost** **12**

**5 Evaluation Metrics** **13**

### 1 Dataset details

#### 1.1 Simulation datasets

##### Overview and Rationale

We benchmarked integration methods on a series of simulated scRNA-seq datasets where the underlying truth is known. These simulated datasets were constructed to include two batches and were designed to reflect three practical challenges often encountered in scRNA-seq data integration: unbalanced batch sizes, rare cell types, and batch-specific cell types. This controlled setting allows for a full characterization of batch effects and enables a robust evaluation of our integration methods.

##### Data Details

We generated synthetic scRNA-seq data using Splatter [1], which employs negative binomial distributions through a hierarchical Gamma-Poisson model. The parameters used were consistent with those applied by Kotliar et al. (2019) [2], estimated from 8000 cells of the organoid dataset in Quadrato et al. (2017) [3].

- **Batches**: 2
- **Cell Types**: 7
- **Number of Cells per Batch**: 2000
- **Number of Genes**: 10,000

Each simulation scenario was repeated ten times using different random seeds to generate a baseline dataset.

##### Scenarios

###### *Unbalanced Batch Sizes*

- **Rationale**: To mimic practical challenges where batch sizes are unequal, which can introduce biases in downstream analyses.
- **Details**:
  - Batch 1 was downsampled to 50% of its original size: **batch1-0.5**.
  - Batch 1 was downsampled to 25% of its original size: **batch1-0.25**.
  - Batch 1 was downsampled to 12.5% of its original size: **batch1-0.125**.

###### *Rare Cell Types*

- **Rationale**: To simulate situations where certain cell types are underrepresented, posing challenges for accurate identification and integration.
- **Details**:
  - Cells labeled as Group1 in each batch were downsampled to 50%: **rare1-0.5**.
  - Cells labeled as Group1 in each batch were downsampled to 20%: **rare1-0.2**.
  - Cells labeled as Group1 in each batch were downsampled to 10%: **rare1-0.1**.

***Batch-Specific Cell Types***

- **Rationale**: To evaluate the performance of integration methods in scenarios where some cell types are not shared across batches, reflecting batch-specific biological conditions.
- **Details**:
  - The number of common cell types was reduced to 5: **common-5**.
  - The number of common cell types was reduced to 3: **common-3**.
  - The number of common cell types was reduced to 1: **common-1**.

For each scenario, three sub-levels were considered, resulting in a total of 9 additional datasets generated based on each baseline dataset. Therefore, there were 10 baseline datasets and 9 modified datasets per baseline, leading to a total of 100 datasets across all sub-scenarios.

##### Previous Use of Data Sets

The same set of parameters and simulation generating are previously used by Wang et al. [4] in their benchmarking. By using these well-characterized datasets, we ensured that our benchmark analysis could be compared with existing studies, thereby validating the robustness and applicability of our integration methods.

#### 1.2 Real dataset

In our study, we applied scCRAFT to integrate nine diverse real datasets, each presenting unique challenges (Table S1). These datasets included small-scale datasets (Bct, Bct_del), commonly used benchmarking datasets (Pancreatic Islets, Lung Atlas, Human Immune datasets), a high-batch dataset (Macaque), and two-species datasets (Lung Two Species, HCL&MCA). These scenarios tested the integration of various techniques and donors, different tissues and species, rare cell types, and large-scale data up to 300,000 cells and 30 batches. The ground truth for these datasets was based on original annotations from the source datasets, consistent with other benchmarking studies.

##### Bct dataset

The Bct dataset comprises mammary epithelial cells collated from three independent studies: Giraddi et al. [5], Pal et al. [6], and Bach et al. [7]. The dataset encompasses cells categorized into three batches—denoted as spk, vis, and wal—each sharing three cell types: basal, luminal_mature, and luminal_progenitor. Post-filtering with minimun 300 genes for cells, the dataset is refined fto include 5,747 cells across 1,222 genes.

The source data for this analysis are retrievable from the Gene Expression Omnibus (GEO) database, with accession codes: GSE111113, GSE98131, GSE103275, and GSE106273. This aggregated dataset, processed and presented by Yu et al. [8], is accessible at https://figshare.com/articles/dataset/Batch_Alignment_of_single-cell_transcriptomics_data_using_Deep_Metric_Learning/20499630/2.

**Table S1** Integration Task Summary

| Integration Task | Cell Number | Batches | Test Features |
|---|---|---|---|
| Bct | 5747 | 3 | Small scale with fewer cell types |
| Bct_del | 4592 | 3 | Small scale with fewer cell types |
| Pancreatic Islet | 16382 | 9 | Technologies, donors, rare cell types |
| Lung | 29256 | 16 | Tissues, laboratories, technologies, heterogeneity in cell composition |
| Human immune | 33506 | 10 | Tissues, laboratories, similar cell types |
| Human immune part | 20292 | 9 | Tissues, laboratories, similar cell types |
| Lung two species | 20760 | 2 | 2-species integration |
| Macaque | 30302 | 30 | Large number of batches |
| HCL&MCA | 372789 | 2 | Large scale, 2-species integration, minimum overlap in cell types |

Note: This table summarizes the integration tasks highlighting various features and challenges of the datasets used in this study.

#### Bct del dataset

The Bct del dataset, derived from the aforementioned Bct dataset, excludes basal cells from the vis and wal batches. This modification results in a focused dataset of 4,592 cells, aimed at enhancing the assessment of integration for partially overlapping cell types. This dataset is also compiled and made available by Yu et al. [8]. It is available for access at the same Figshare link provided above.

#### Human Immune dataset

The Human Immune dataset encompasses cells from ten human samples across two distinct tissues: bone marrow and peripheral blood. Bone marrow samples were sourced from the study conducted by Oetjen et al. [9], whereas peripheral blood samples were compiled from multiple sources, including publicly available data from 10x Genomics [10], Freytag et al. [11], Sun et al. [12], and Villani et al. [13]. Comprehensive details on dataset procurement, the varied experimental protocols employed, and the criteria for sample selection are detailed in the work by Luecken et al. [14]. The datasets leveraged in this analysis are accessible via the GEO database, with accession codes GSE120221, GSE107727, GSE115189, GSE128066, and GSE94820, along with the 10X Genomics website (PBMC10k: https://support.10xgenomics.com/single-cell-gene-expression/datasets/3.0.0/pbmc_10k_v3) [14] for the PBMC10k dataset. Following rigorous quality control measures, the dataset was distilled to include 33,506 cells across 12,303 genes, spanning ten different batches.

#### Human Immune part dataset

The Human Immune part dataset is a refined version of the widely used Human Immune dataset, excluding data from Villani et al. [13], which were exclusively based on TPM values. This exclusion results in a refined compilation of 32,259 cells. This dataset, processed and made available by Lotfollahi et al. [15], can be accessed via https://github.com/theislab/scarches-reproducibility?tab=readme-ov-file. Including this refined dataset, in addition to the full Human Immune dataset, allows us to demonstrate the robustness of our method across different quality control levels. The refined dataset, which has been used for benchmarking by Lotfollahi et al. [15], underscores its relevance and applicability in advanced integration method benchmarking.

#### Pancreatic Islet dataset

Pancreatic Islet dataset incorporates five publicly available pancreatic islet datasets [16–20], aggregating to a comprehensive collection of 16,382 cells. The detailed compilation process, including criteria for cell selection and quality control, as well as the experimental protocols for each batch, is thoroughly documented by Luecken et al. [14], ensuring transparency and reproducibility of our analysis.

The datasets are archived within the Gene Expression Omnibus (GEO) database, accessible under accession codes GSE81076, GSE85241, GSE86469, GSE84133, and GSE81608. Additionally, data are also retrieved from the ArrayExpress database with the accession code "E-MTAB-5061 [https://www.ebi.ac.uk/biostudies/arrayexpress/studies/E-MTAB-5061]", as detailed in the benchmarking study by Luecken et al [14].

#### Lung atlas dataset

The lung atlas is sourced from the study by Vieira Braga et al. [21], and is hosted within the Gene Expression Omnibus (GEO) database, with the accession code GSE130148. This dataset represents a valuable resource for exploring the cellular landscape of the lung. The process involved in dataset compilation, including the quality control, the experimental protocols used for each batch, and specific details regarding the acquisition of these datasets, is detailed outlined in the benchmarking study provided by Luecken et al [14].

#### Lung two species dataset

The Lung Two Species dataset is designed to facilitate cross-species comparisons, comprising lung cells from both mouse and human, segregated into two distinct batches. The original dataset is available through the Gene Expression Omnibus (GEO) database under the accession code GSE133747, as reported by Raredon et al. [22]. This dataset was further processed and compiled by Yu et al. [8], resulting in a refined version containing a total of 20,760 cells and 62,781 genes, which is accessible at https://figshare.com/articles/dataset/Batch_Alignment_of_single-cell_transcriptomics_data_using_Deep_Metric_Learning/20499630/2.

#### Macaque Retina dataset

The Macaque Retina dataset, generated by Peng et al. [23], is a dataset with clearly defined cell types for exploring retinal cell compositions in macaques with each sample defined as one batch for integration. Hosted in the Gene Expression Omnibus (GEO) database with the accession code GSE118480. Subsequently processed and made accessible by Yu et al. [8], the dataset can be found at https://figshare.com/articles/dataset/Batch_Alignment_of_single-cell_transcriptomics_data_using_Deep_Metric_Learning/20499630/2

#### HCL-MCA dataset

The Human Cell Landscape (HCL) dataset and the Mouse Cell Atlas (MCA) dataset serve as foundational elements for cross-species comparative analysis, with both datasets retrieved from https://figshare.com/articles/HCL_DGE_Data/7235471. They were aggregated by Lotfollahi et al. [15] and made accessible at https://github.com/theislab/scarches-reproducibility?tab=readme-ov-file. The merged collection comprising 249,845 cells from 63 cell types in the HCL batch and 122,944 cells from 50 distinct cell types in the MCA batch after the filtering.

### 2 Benchmarking Methods

In our benchmarking study, we evaluated several computational methods for single-cell data integration, focusing on their ability to balance batch correction with the preservation of biological variation. These methods fall into two main categories: reference-based and mixing-loss-based integration.

**reference-based methods**, such as BBKNN, Scanorama, and Seurat, identify reference cells across batches to align datasets. These approaches are effective but depend on the availability of high-quality references.

**Mixing-loss-based methods**, including Harmony and scVI, use a combination of reconstruction and batch-mixing losses to integrate data. While powerful, these methods can sometimes over-correct or under-correct batch effects. Hybrid approaches like iMAP and scDML incorporate aspects of both strategies for improved stability.

A summary of the methods used, including version information, specific settings, and tutorial references, is provided in Table S2.

**Table S2** Summary of Benchmarking Methods

| Method | Specific Settings | Tutorial Website |
|---|---|---|
| BBKNN (v1.6.0) | Default setting with k = 3 neighbors for intra-batch comparisons | https://bbknn.readthedocs.io/en/latest/ |
| Scanorama | scanorama.correct_scanpy with return_dimred=True option | https://github.com/brianhie/scanorama |
| Seurat (v5) | Used RPCAIntegration with SCTransform normalization; top 50 PCA embeddings | https://satijalab.org/seurat/ |
| Harmony | Default settings; PCA space initialization | https://github.com/slowkow/harmonypy |
| scVI | 50-dimensional latent space, 128 nodes per hidden layer, 2 layers | https://docs.scvi-tools.org/en/stable/tutorials/notebooks/scrna/harmonization.html |
| iMAP | 50 training epochs for initial encoding, 40 epochs for GAN-based refinement | https://github.com/Svvord/iMAP/blob/master/tutorials/cell_lines_tutorial.md |
| scDML | 50-dimensional embedding; number of clusters (n_cluster) set to match true cell types | https://github.com/eleozzr/scDML/blob/main/tutorial/tutorial2.ipynb |

* Note: All methods except for BBKNN were configured to output a 50-dimensional embedding.

### 3 Detailed algorithm

Table S3 provides a detailed breakdown of the layers and operations involved in each component of the scCRAFT model, including the encoder, decoder, and discriminator networks.

**Table S3** scCRAFT Model Neural Network Architecture

| Name | Operation | NoF/Kernel Dim. | Activation | Input |
|---|---|---|---|---|
| | | **Encoder** | | |
| Layer_1 | FC | 1024 | ReLU | data |
| Layer_2 | FC | 512 | ReLU | Layer_1 |
| mean | FC | 256 | Linear | Layer_2 |
| var | FC | 256 | Linear | Layer_2 |
| latent | Multinormal | 256 | - | [mean, var] |
| | | **Decoder** | | |
| **Generator 1** | | | | |
| Layer_11 | FC | 512 | ReLU | [latent, condition] |
| Layer_12 | FC | 1024 | ReLU | Layer_11 |
| **Generator 2** | | | | |
| Layer_21 | FC | 512 | ReLU | condition |
| Layer_22 | FC | 1024 | ReLU | Layer_21 |
| **Combine** | | | | |
| combine | ReLU | p_dim | - | [Layer_12, Layer_22] |
| px_scale | FC | p_dim | - | combine |
| px_r | FC | p_dim | - | combine |
| | | **Discriminator** | | |
| Layer_1 | FC | 128 | ReLU | latent |
| Layer_2 | FC | 128 | ReLU | Layer_1 |
| classification | FC | domain_number | Softmax | Layer_2 |

Note: FC stands for Fully Connected. NoF refers to Number of Features.

**Hyperparameters:**

Loss: NB

Optimizer: Adam

Learning Rate: 0.001

$\epsilon$: 0.01

Batch Size: 1024

Algorithm 1 details the steps to compute the triplet loss within a mini-batch, ensuring that embeddings for similar cell types are closer together while separating different cell types. It starts by counting the occurrences of each label in every batch (line 2). Then, it initializes an empty list to store calculated triplet losses (line 3). As it loops through each batch and low-resolution label (lines 4-5), the algorithm identifies the indices of cells within the same low-resolution label (connected) and those with different low-resolution labels (not-connected) within one batch (lines 6-7). It then generates pairs of connected examples and samples corresponding not-connected examples (lines 8-9). For each connected pair, the embeddings for the reference, connected, and not-connected cells are retrieved (lines 14-15). The algorithm checks whether the high-resolution labels of the reference and connected cells match to determine whether this is a reference, positive and negative triplet (line 17) and finally computes the triplet loss based on the distance between reference-positive and reference-negative pairs (lines 18-19).

**Algorithm 1** Triplet Loss Computation with Connected and Non-Connected Pairs

**Require:** $embeddings(z)$, $l_{\text{low}}$, $l_{\text{high}}$, $batch\_ids(v)$
**Require:** $margin = 5.0$, $num\_triplets\_per\_label = 15$
**Ensure:** $average\_triplet\_loss$

1: Initialize all parameters.
2: $label\_counts\_per\_batch \leftarrow$ CountLabelsPerBatch($labels$, $batch\_ids$)
3: $triplets \leftarrow []$
4: **for** $v$, $(unique\_low\_labels, counts)$ in $label\_counts\_per\_batch$ **do**
5:     **for** $label$ in $unique\_low\_labels$ **do**
6:         $connected\_indices \leftarrow l_{\text{low}} = label \wedge v = batch\_id$
7:         $non\_connected\_indices \leftarrow l_{\text{low}} \neq label \wedge v = batch\_id$
8:         $connected\_pairs \leftarrow$ Combinations($connected\_indices$, 2)
9:         $sampled\_connected\_pairs \leftarrow$ Sample($connected\_pairs$, $num\_triplets\_per\_label$)
10:         $sampled\_non\_connected\_indices \leftarrow$ Sample($non\_connected\_indices$, $|\text{sampled\_connected\_pairs}|$)
11:         **for** $i$ in $range(0, |\text{sampled\_connected\_pairs}|)$ **do**
12:             $(reference\_idx, connected\_idx) \leftarrow sampled\_connected\_pairs[i]$
13:             $non\_connected\_idx \leftarrow sampled\_non\_connected\_indices[i]$
14:             $reference \leftarrow z[reference\_idx]$
15:             $connected \leftarrow z[connected\_idx]$
16:             $non\_connected \leftarrow z[non\_connected\_idx]$
17:             **if** $l_{\text{high}}[reference\_idx] = l_{\text{high}}[connected\_idx]$ **then**
18:                 $positive = connected$
19:                 $negative = non\_connected$
20:                 $triplet\_loss \leftarrow \max(0, ||reference - positive||_2$
21:                     $-||reference - negative||_2 + margin)$
22:                 $triplets.append(triplet\_loss)$
23:             **end if**
24:         **end for**
25:     **end for**
26: **end for**
27: **if** $triplets$ not empty **then**
28:     **return** mean($triplets$)
29: **else**
30:     **return** 0
31: **end if**

Algorithm 2 outlines the comprehensive training procedure for the scCRAFT model. The process begins with preprocessing steps (lines 2-3) that involve clustering to obtain low and high-resolution labels, followed by initializing necessary parameters. During training (lines 4-23), the algorithm iteratively processes batches of data, where it first loads and sorts the data for domain alignment (lines 6-7). It then computes the latent space embeddings (line 8) and calculates the triplet loss (line 9) to ensure that similar cell types are closer together in the embedding space. The decoder reconstructs the data (line 10), and the algorithm computes the ELBO and cosine similarity losses (line 11). The discriminator is optimized over several iterations (lines 12-14), and the adversarial loss is computed (line 15). Finally, the total loss, combining all components, is calculated (line 16) and used to update the encoder and decoder (line 17). The process repeats across epochs, and the trained encoder and decoder modules are returned at the end (line 21).

**Algorithm 2** scCRAFT Training Procedure with Discriminator and Triplet Loss

**Require:** AnnData object *adata*, batch key *batch_key*, dimension of latent space *z_dim*

**Ensure:** Trained model components: Encoder *EC*, Decoder *Dec*

1: **Preprocessing:** Execute dual-resolution clustering to obtain low and high resolution labels: $l_{\text{low}}, l_{\text{high}}$
2: Initialize with *p_dim*, *v_dim*, and *z_dim*
3: **for** $epoch = 1$ to $epochs$ **do**
4:     Load *adata* with balanced batches based on *batch_key*
5:     Activate training mode for $EC$, $Dec$, and $D_Z$
6:     **for all** batches $(x, v, l_{\text{low}}, l_{\text{high}})$ from the data loader **do**
7:         Sort $(x, v, l_{\text{low}}, l_{\text{high}})$ by $v$ for domain alignment
8:         $z \leftarrow EC(x)$
9:         Calculate triplet loss $L_{triplet}$ based on $z$, $l_{\text{low}}$, $l_{\text{high}}$, and $v$
10:         $\tilde{x} \leftarrow Dec(z, v)$
11:         Compute ELBO loss $L_{\text{ELBO}}$ and cosine similarity loss $L_{cos}$
12:         **for** $k = 1$ to 10 iterations **do**
13:             Optimize $D_Z$ with $v$
14:         **end for**
15:         Calculate adversarial loss $L_{adv}$ for generator against $D_Z$
16:         $L_{total} \leftarrow L_{\text{ELBO}} + L_{cos} + L_{triplet} - L_{adv}$
17:         Update $EC$ and $Dec$ using their optimizer
18:     **end for**
19: **end for**
    **return** $EC$, $Dec$ as the trained model components

### 4 Additional discussion on computational cost

In our benchmarking study of deep learning methods for single-cell data analysis, we observed different trends in computational efficiency across different models, particularly in how training time correlates with dataset characteristics. Notably, the training time for some models remained relatively stable across varying dataset sizes, while others exhibited a linear increase in computational time as the dataset size grew.

These observed trends can be explained by the sampling schemes employed by each model. For example, **scVI** utilizes an early stopping mechanism capped at 400 epochs and pre-defines the number of epochs based on the cell count. This approach effectively decouples the training duration from the total dataset size, allowing scVI to maintain consistent training times regardless of how large the dataset is.

In contrast, **iMAP** introduces a distinct training strategy that begins by sampling an equal and fixed number of cells from each batch, followed by a mix-up procedure to create the final mini-batches for training within each epoch. As a result, the training time for iMAP is influenced more by the number of batches rather than the total cell count. An increase in the number of batches extends the matching steps, making the model's computational time more dependent on batch configuration than on overall dataset size.

**scDML**, on the other hand, sets its training epochs to a fixed number, which results in a training duration that scales linearly with the number of cells. This direct link between computational time and dataset size explains the linear increase in training time observed with larger datasets.

**scCRAFT** employs a similar batch-wise sampling method like iMAP by random selection of 512 cells from each batch to create a pool, followed by the random sampling of 1024 cells per mini-batch from this pool. Consequently, when the cell count per batch exceeds 512, the training time for scCRAFT is primarily influenced by the number of batches rather than the total number of cells. We predefine the total training epochs to be 150. For datasets with a low number of batches but a high cell count, an additional adjustment is made: if the cell count exceeds 100,000, the number of epochs is calculated as $\text{int}(1.5 \times \text{number of cells}/(\text{number of batches} \times 512))$. If this calculated number exceeds 150, it is used; otherwise, the epoch count remains at 150. This adjustment accounts for the increase in training time observed when the cell numbers exceed 100,000, explaining the model's time complexity in such cases.

### 5 Evaluation Metrics

#### 5.1 Overview

To evaluate single-cell data integration methods, we focused on two main categories: (1) biological variance conservation and (2) batch effect correction. The biological variance conservation metrics assess how well the methods preserve the original biological signals. The batch correction metrics evaluate the effectiveness of methods in minimizing batch effects. Composite scores takes the average of metrics in each categories to obtain a more comprehensive scores. The Overall Score integrates these two aspects, with a weighting scheme that reflects their relative importance in single-cell data integration. Supplementary metrics work as additional assessment of different integration methods performance.

Table S4 summarizes the key metrics used in our evaluation, including the composite scores that combine these metrics into a single value. The composite scores are bolded to indicate their significance in the overall evaluation.

#### 5.2 Details on Biological Conservation Metrics:

For calculating the cluster based biological conservation metrics – NMI and ARI, clustering of the integrated data is essential. Suggested and implemented by Luecken et al. [14], we employ the Louvain clustering algorithmand select a resolution that optimizes the Normalized Mutual Information (NMI) value. This optimization of Louvain clustering across a defined resolution range provides a more accurate reflection of biological conservation for these two metrics:

1. **Normalized Mutual Information (NMI):** $NMI$ is utilized to quantify the correspondence between cell-type labels and clusters derived from the integrated dataset through Louvain clustering. By scaling the overlap using the mean of the entropy terms for both cell-type and cluster labels, $NMI$ offers a normalized measure ranging from 0 (no correlation) to 1 (perfect correlation), ensuring a relative assessment of cluster purity and label consistency.
2. **Adjusted Rand Index (ARI):** $ARI$ measures the congruence between the cell-type labels and the clusters identified by optimized Louvain clustering [24]. By adjusting for the chance grouping of labels, $ARI$ provides a scale from 0 (random labeling) to 1 (perfect match), offering insights into the clustering efficiency and the preservation of cell identity post-integration [25].
3. **Cell Type Average Silhouette Width (ASW):** $ASW$ evaluates the clarity of separation and cohesion within and between clusters, respectively.

$$ASW = \frac{1}{N}\sum_{i=1}^{N}\frac{b(i)-a(i)}{\max\{a(i),b(i)\}}$$

For each cell $i$, $a(i)$ is the mean distance to the other cells in the same cell type, and $b(i)$ is the mean distance to the nearest cluster that $i$ is not a part of. $cASW = (ASW+1)/2$ is the final score. By calculating the silhouette width for each cell and averaging these values, $ASW$ presents a metric that discerns the degree of cluster

**Table S4** Summary of Evaluation Metrics

| **Metric** | **Description** |
|---|---|
| **Biological Variance Conservation** | |
| Normalized Mutual Information (NMI) | Measures overlap between cell-type labels and clusters. |
| Adjusted Rand Index (ARI) | Assesses clustering accuracy relative to known labels. |
| Cell Type Average Silhouette Width (ASW_cell) | Evaluates cluster cohesion and separation. |
| Cell Type Local Inverse Simpson's Index (cLISI) | Measures cell type diversity within local neighborhoods. |
| **Batch Effect Correction** | |
| Average Silhouette Width for Batches (ASW_batch) | Measures distance between different batches. |
| Principal Component Regression (PCR_batch) | Quantifies batch-related variance within principal components. |
| Graph Connectivity | Evaluates cohesiveness of cells of the same type within the kNN graph. |
| k-Nearest Neighbor Batch Effect Test (kBET) | Assesses local neighborhood consistency with global label distribution. |
| Batch Label Identity Score Index (bLISI) | Measures batch diversity within local neighborhoods. |
| Proportion of True Positive Cells | Assesses the proportion of positive cells which mirror the global batch distribution. |
| **Supplementary Metrics** | |
| LISI F1 Score | Combines cLISI and bLISI to measure overall integration quality. |
| **Composite Scores** | |
| **Biological Conservation Score** | Average of NMI, ARI, ASW_cell, and (1-cLISI). |
| **Batch Mixing Score** | Average of bLISI, ASW_batch, kBET, Graph Connectivity, PCR_batch, and true positive rate. |
| **Overall Score** | Weighted sum of Biological Conservation Score (60%) and Batch Mixing Score (40%). |

overlap and misclassification, with values ranging from 0 (poor clustering) to 1 (distinct and dense clustering). This measure is pivotal in assessing the method's ability to maintain distinct biological entities within the integrated dataset.

4. **Cell Type Local Inverse Simpson's Index (cLISI):** *cLISI* [26] is a metric designed to measure the diversity of cell types within local neighborhoods, effectively quantifying the degree of biological variance conservation.

$$cLISI = 1 - \frac{\sum_{c=1}^{C} p(c)^2 / C - 1}{C - 1}$$

$C$ represents the number of unique cell types, and $p(c)$ is the probability of cell type c within a local neighborhood (default 30 cells). The final cLISI score for a dataset is then obtained by averaging the cLISI values across all cells in the dataset. By calculating the inverse Simpson's Index based on cell type probabilities within local neighborhoods, *cLISI* offers a scale indicative of the effective number of cell types present. This metric, after min-max scaling, is instrumental in evaluating the local mixing of cell types, with an ideal goal of maximizing diversity while retaining biological specificity.

#### 5.3 Details on Batch correction metrics:

1. **Average Silhouette Width for Batches (ASW_batch):** The *ASW_batch* for each cell $i$ is calculated as:

$$\text{ASW\_batch} = \frac{1}{N}\sum_{i=1}^{N}\frac{b(i)-a(i)}{\max\{a(i),b(i)\}}$$

where: - $a(i)$ is the average distance from cell $i$ to all other cells in the same batch, - $b(i)$ is the average distance from cell $i$ to cells in the nearest different batch.

The final *ASW_batch* score is then transformed as follows:

$$\text{ASW\_batch\_final} = 1 - |\text{ASW\_batch}|$$

here a score of 1 indicates perfect batch mixing, with no discernible batch effects, while a score of 0 indicates poor mixing, where cells remain clustered by batch rather than by biological similarity.

2. **Principal Component Regression (PCR) for Batch Effects:** *PCR_batch* [27] quantifies the variance attributed to batch effects within each principal component ($PC_k$). Let $M$ represent the data matrix, where rows correspond to cells and columns to genes, and $K$ denote the total number of principal components considered in the analysis. The variance attributed to batch $b$ within each principal component $\text{PC}_k$ is calculated as:

$$\text{Var}(M|b) = \sum_{k=1}^{K}\text{Var}(M|\text{PC}_k)\times R^2(\text{PC}_k|b)$$

Here $\text{Var}(M|\text{PC}_k)$ is the variance in the data explained by the $k$-th principal component. $R^2(\text{PC}_k|b)$ is the coefficient of determination from a linear regression model where the batch labels are treated as independent variables, and the principal component scores for $\text{PC}_k$ are the dependent variable.

This approach measures the impact of batch effects on the dataset's variance, identifying methods that proficiently reduce batch-related variance.

3. **Graph Connectivity:** This metric assesses the cohesiveness of cells from the same cell type within the k-nearest neighbor (kNN) graph.

$$GC = \frac{1}{|C|}\sum_{c\in C}\frac{|\text{LCC}(G(N_c,E_c))|}{N_c}$$

where $C$ represents the set of all cell types. For each cell type $c$, $G(N_c, E_c)$ denotes the kNN subgraph constructed exclusively for that cell type $c$. $N_c$ is the number of cells of type $c$. $E_c$ represents the edges in the kNN graph that connect only cells of type $c$. $|\mathrm{LCC}(G(N_c, E_c))|$ refers to the number of cells in the largest connected component of this subgraph.

A higher $GC$ score suggests a well-mixed integration, indicating that cells sharing the same type label are directly interconnected across different batches.

4. **k-Nearest Neighbor Batch Effect Test (kBET):** The *kBET* metric[27] assesses the effectiveness of batch effect correction by evaluating whether the batch composition within local neighborhoods matches the global batch distribution after data integration. It works as follows: A k-nearest neighbor (kNN) graph is constructed for each cell, typically with $k = 50$, to define its local neighborhood. The test compares the batch label distribution within each neighborhood to the global batch distribution across the entire dataset. A statistical test is performed to determine if the local batch composition deviates significantly from the global expectation. Neighborhoods that show significant deviation are "rejected," indicating poor batch mixing.

   In cases where the k-nearest neighbor (kNN) graph is disconnected, kBET is performed on each connected component, with neighborhood sizes adapted to the specific cell distribution within each batch, typically ranging from 10 to 100 neighbors. If more than 25% of cells fall within components too small for kBET analysis (fewer than $k \times 3$ neighbors), a default kBET score of 1 is assigned, indicating insufficient batch correction. The final kBET score is calculated by averaging the individual batch scores and inverting them by subtracting from 1, where a score close to 1 signifies effective batch mixing, and a score close to 0 indicates poor mixing.

5. **Proportion of Positive and True Positive Cells:** This metric[28] evaluates integration quality at the single-cell level by categorizing cells as positive or negative based on their neighborhood composition. Positive cells are those surrounded by the same cell type; Cells surrounded by the same cell type are labeled as positive; otherwise, they are negative. Among the positive cells, a cell is further classified as a true positive if the batch distribution in its neighbor matches the global batch distribution. The proportion of true positive is defined as the number of true positive cells divided by total cell number, which measures the effectiveness of batch mixing. In this benchmarking, we applied true positive rate as a metrics for batch correction evaluation and positive rate as a supplementary metrics.

6. **Batch Label Identity Score Index (bLISI):** *bLISI* [4] is a metric designed to measure the diversity of batch labels within local neighborhoods, effectively quantifying the degree of batch mixing.

$$bLISI = \frac{\sum_{b=1}^{B} p(b)^2 / B - 1}{B - 1}$$

   Here, $B$ represents the number of unique batches, and $p(b)$ is the probability of batch $b$ within a local neighborhood (default 30 cells). By calculating the inverse Simpson's Index based on batch label probabilities within local neighborhoods, *bLISI* offers a scale indicative of the effective number of batches present.

This metric, after min-max scaling, is instrumental in evaluating the local mixing of batches, with an ideal goal of maximizing batch homogeneity while ensuring that biological variance is retained.

#### 5.4 Details on Composite Scores

We have devised composite scoring metrics to enable a comprehensive and comparative analysis of data integration methods, encompassing both biological conservation and batch correction effectiveness. These composite scores are constructed by first scaling individual metrics through min-max normalization across competing methods to ensure a fair evaluation. The scaled metrics are then averaged within their respective categories to yield the composite scores.

The composite scores are constructed as follows:

1. **Biological Conservation Score:** This score is the mean of the first four metrics that are indicative of biological signal preservation. The formula is given by:

$$\text{Biological Conservation Score} = \frac{\text{ASW}_{\text{label}} + \text{ARI} + \text{NMI} + (1 - \text{cLISI})}{4}$$

2. **Batch Mixing Score:** This score is the average of the metrics that reflect the extent of batch effect correction. It is computed as:

$$\text{Batch Mixing Score} = \frac{\text{bLISI} + \text{ASW}_{\text{batch}} + \text{kBET Accept Rate} + \text{graph connectivity} + \text{PCR}_{\text{batch}} + \text{true pos rate}}{6}$$

3. **Overall Score:** The final score combines the biological conservation and batch mixing scores, weighted to reflect their respective importance[14]. The overall score is calculated as:

$$\text{Overall Score} = 0.6 \times \text{Biological Conservation Score} + 0.4 \times \text{Batch Mixing Score}$$

The Biological Conservation Score captures the method's ability to maintain biologically relevant information, while the Batch Mixing Score assesses the effectiveness of batch effect removal. The Overall Score, a weighted sum of these two scores, provides a singular metric of a method's performance across both biological conservation and batch mixing.

#### 5.5 Details on Supplementary Integration Metrics:

Except the above biological conservation and batch mixing metrics used for direct benchmarking, we also include one one overall evaluation metrics $LISI_{F1}$ which is calculated by:

$$\text{LISI}_{F1} = (2 * \text{cLISI} * \text{bLISI})/(\text{cLISI} + \text{bLISI})$$

to aid the benchmarking.

### 6 Supplementary Figures

#### 6.1 UMAP and metrices evaluation for simulation and real datasets

The arrangement of panels is the same across Fig. S1-16, and we provide their shared Figure legends here to avoid redundancy:

(a) UMAP visualization contrasting the unintegrated dataset with its integrated counterparts using scCRAFT and two other most popular benchmarking methods.

(b) Scatterplot of average bio-conservation score against the average batch correction score for each method.

(c) details the specific values and rankings for each evaluation metric across the different methods. The biological and batch correction scores are obtained through aver- aging from metrics corresponding to their respective domains. The overall score is calculated in the same way as the above. The diameter of each circle is determined through a linear rescaling of its corresponding values to a range between 0.1 and 1. Conversely, the length of each bar representing the average score is directly proportional to its actual value. Methods are analyzed and ranked according to their composite achievements in both biological conservation and batch correction.

For Fig. S17-19, some panels have appeared in the main text and only new ones are remained: For Fig. S17 and Fig. S18, previous panel (b) is not included, and for Fig. S19, only panel (a) is kept.

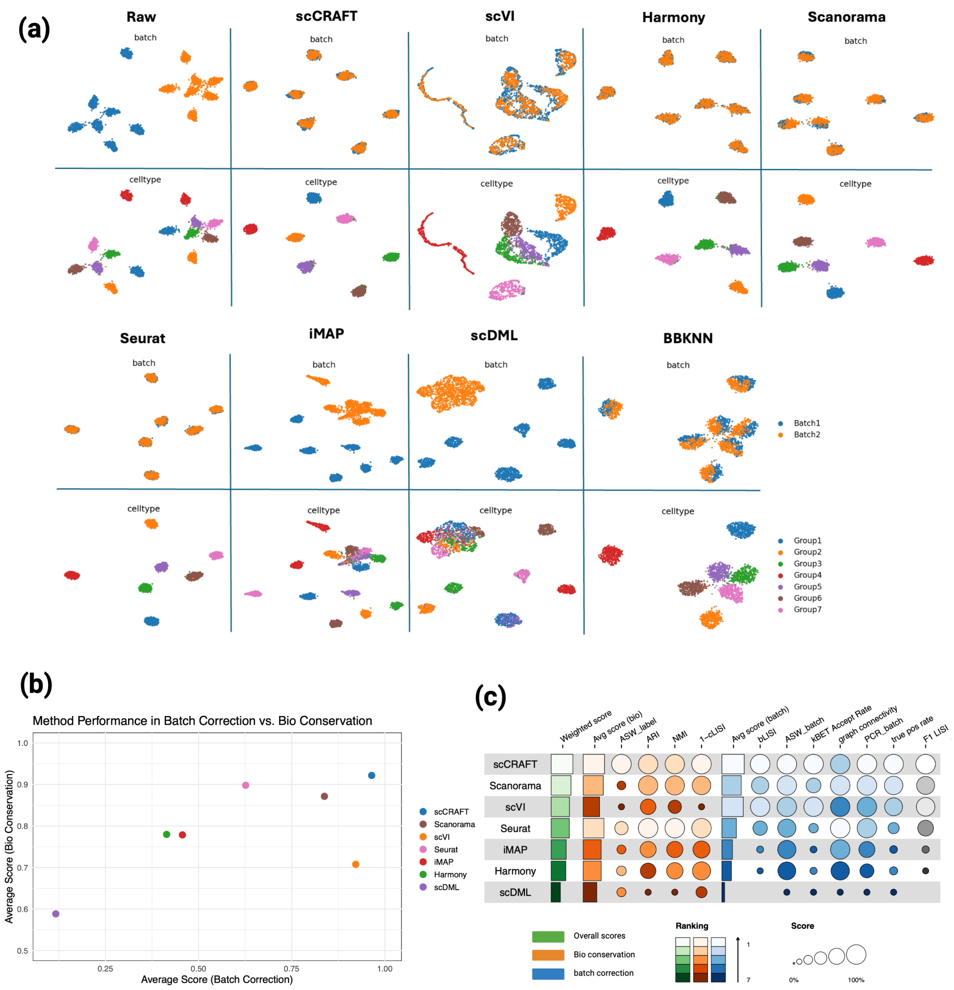

**Fig. S1** Benchmarking Results on simulation task: Baseline. See Suppl Section 6 for details.

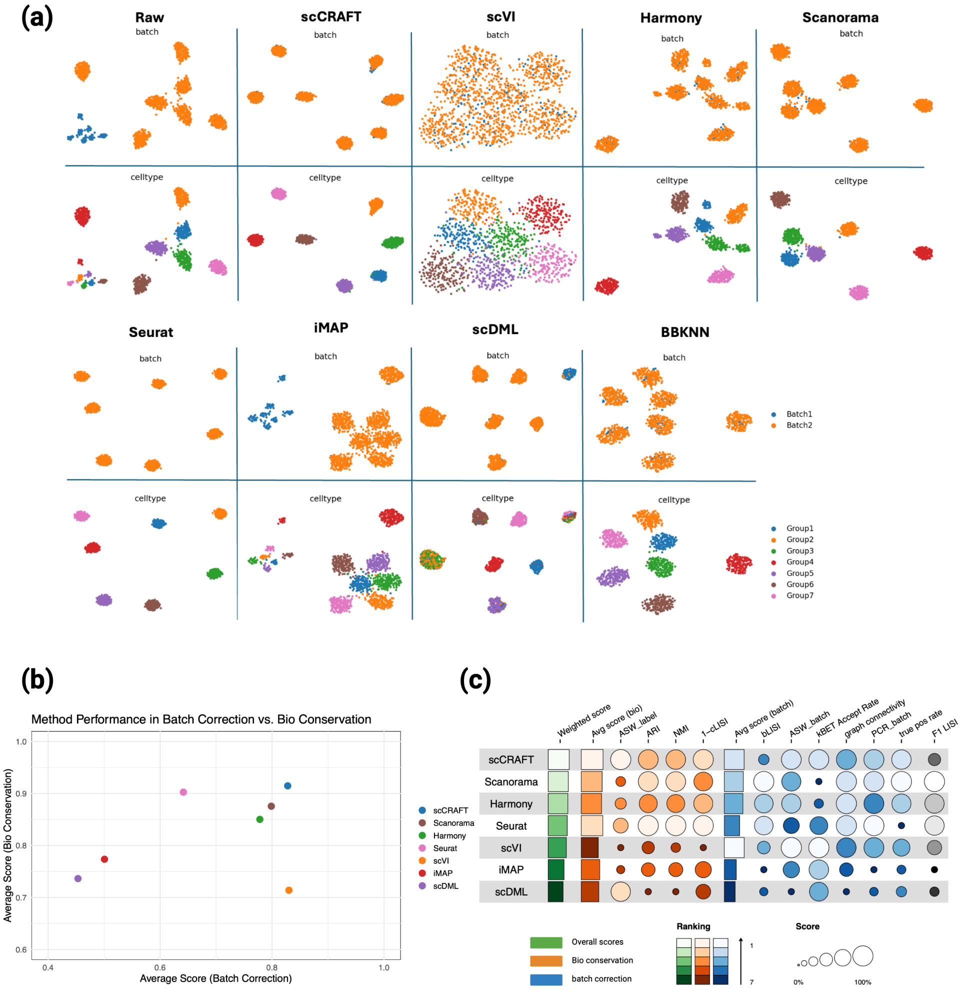

**Fig. S2** Benchmarking Results on simulation task: Batch 0.125. See Suppl Section 6 for details.

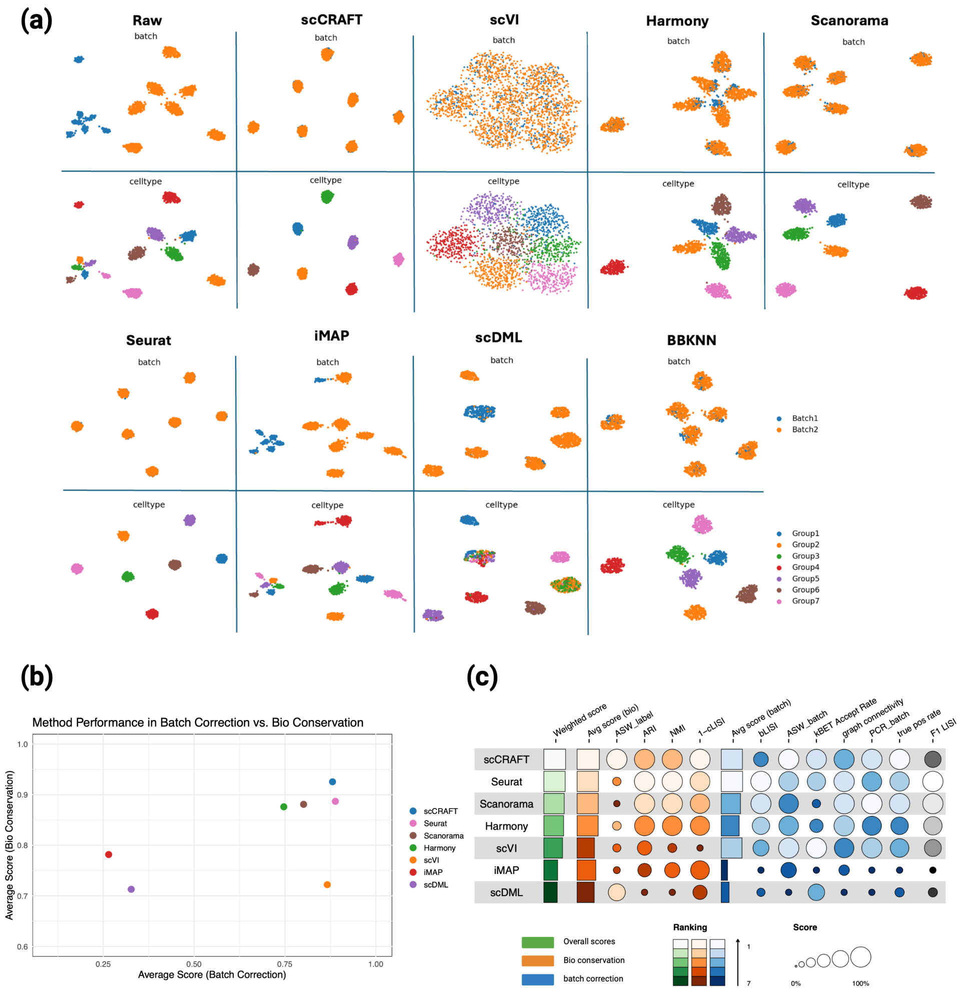

**Fig. S3** Benchmarking Results on simulation task: Batch 0.250. See Suppl Section 6 for details.

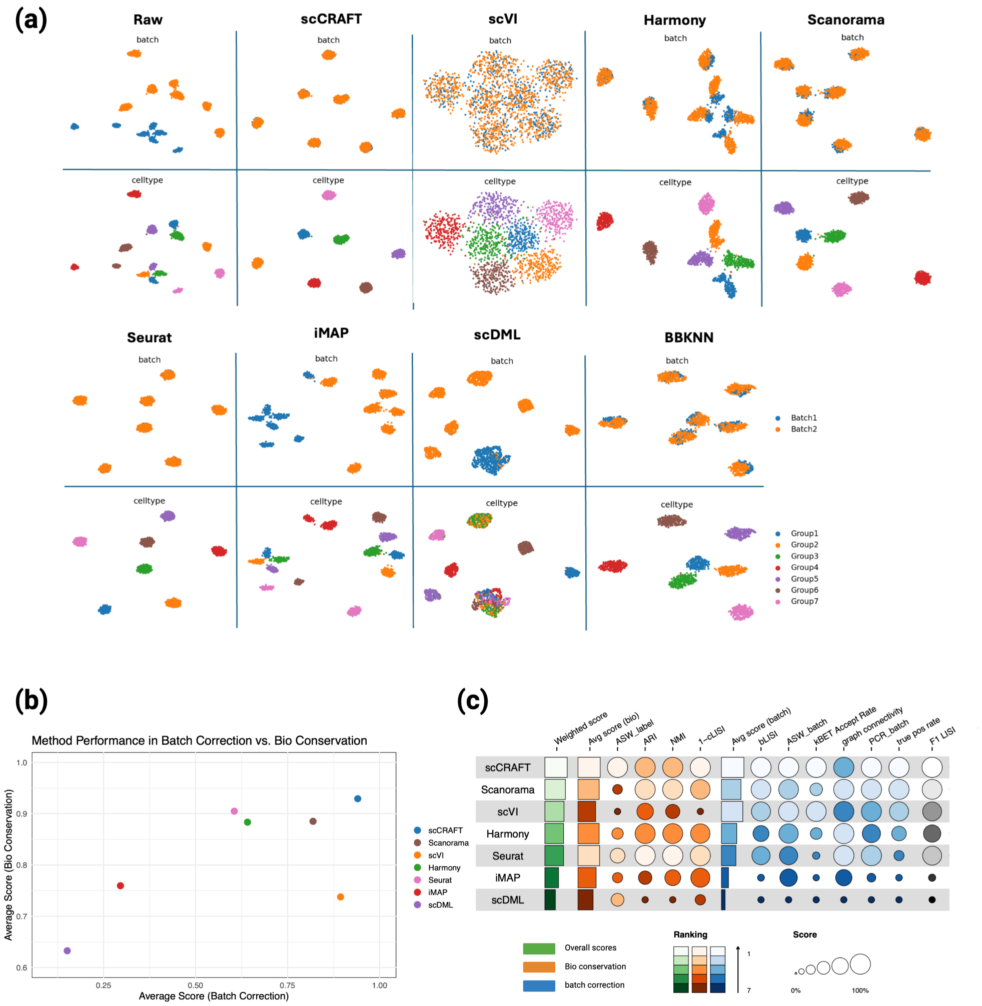

**Fig. S4** Benchmarking Results on simulation task: Batch 0.500. See Suppl Section 6 for details.

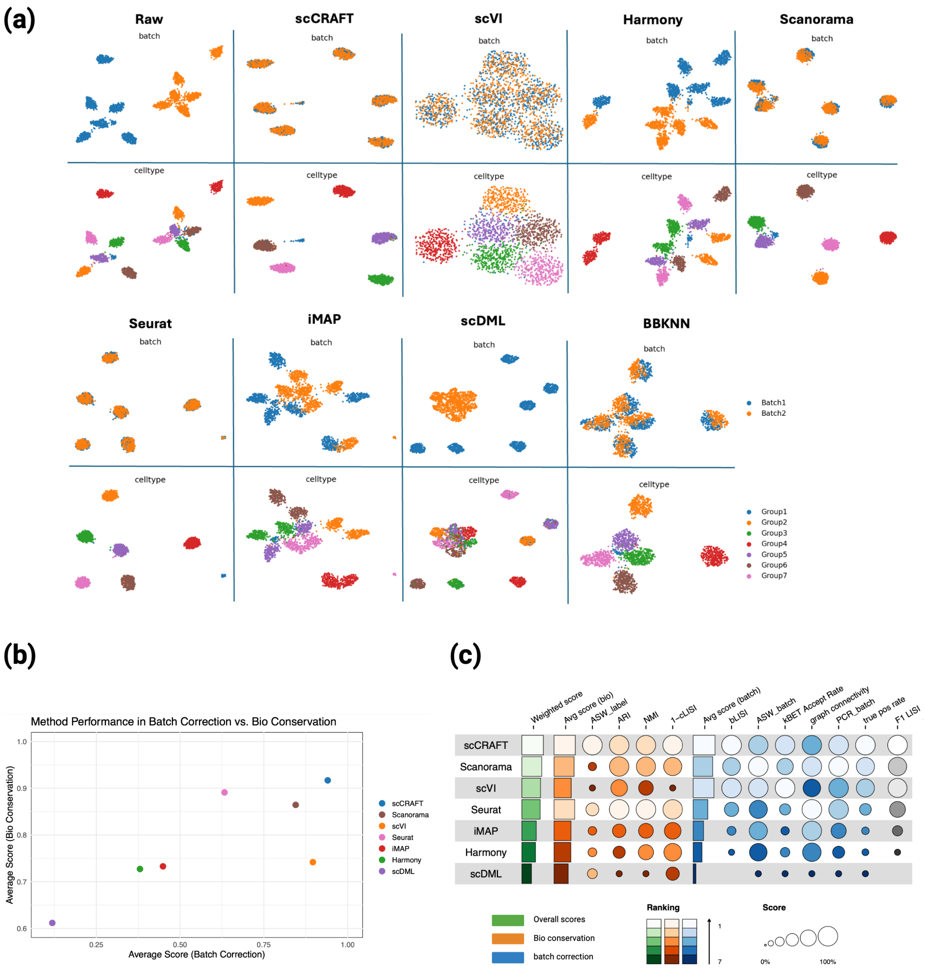

**Fig. S5** Benchmarking Results on simulation task: Rare 0.1. See Suppl Section 6 for details.

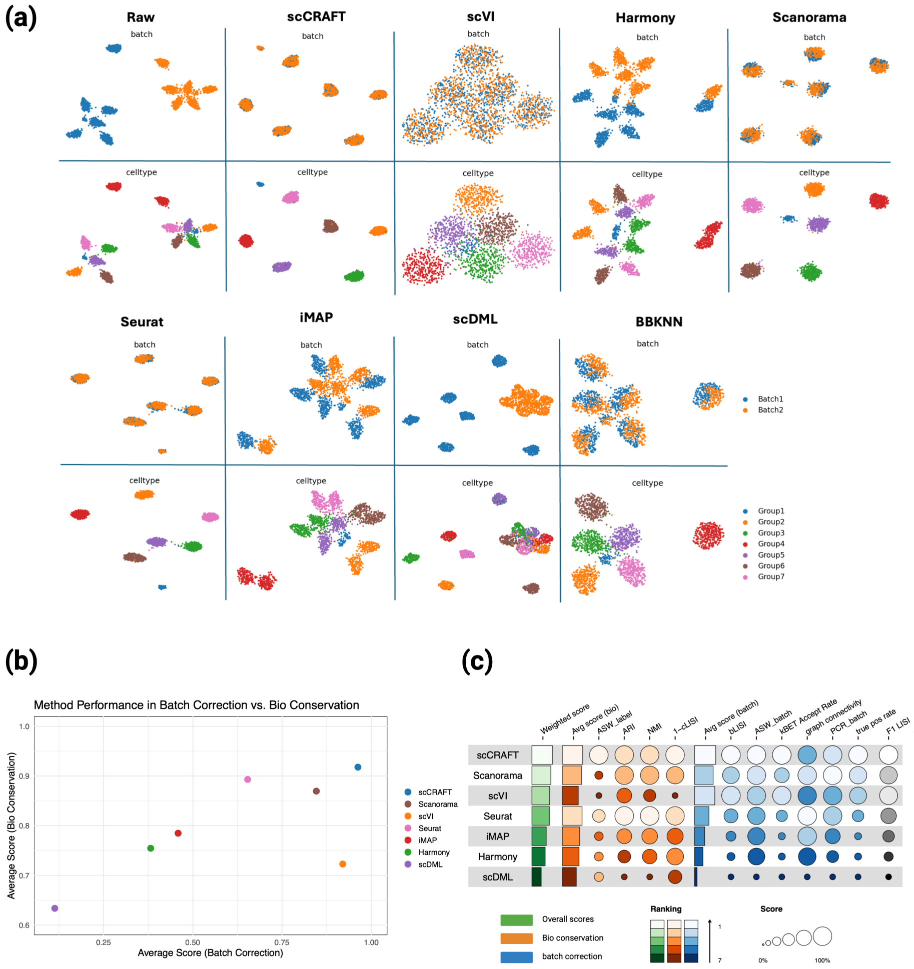

**Fig. S6** Benchmarking Results on simulation task: Rare 0.2. See Suppl Section 6 for details.

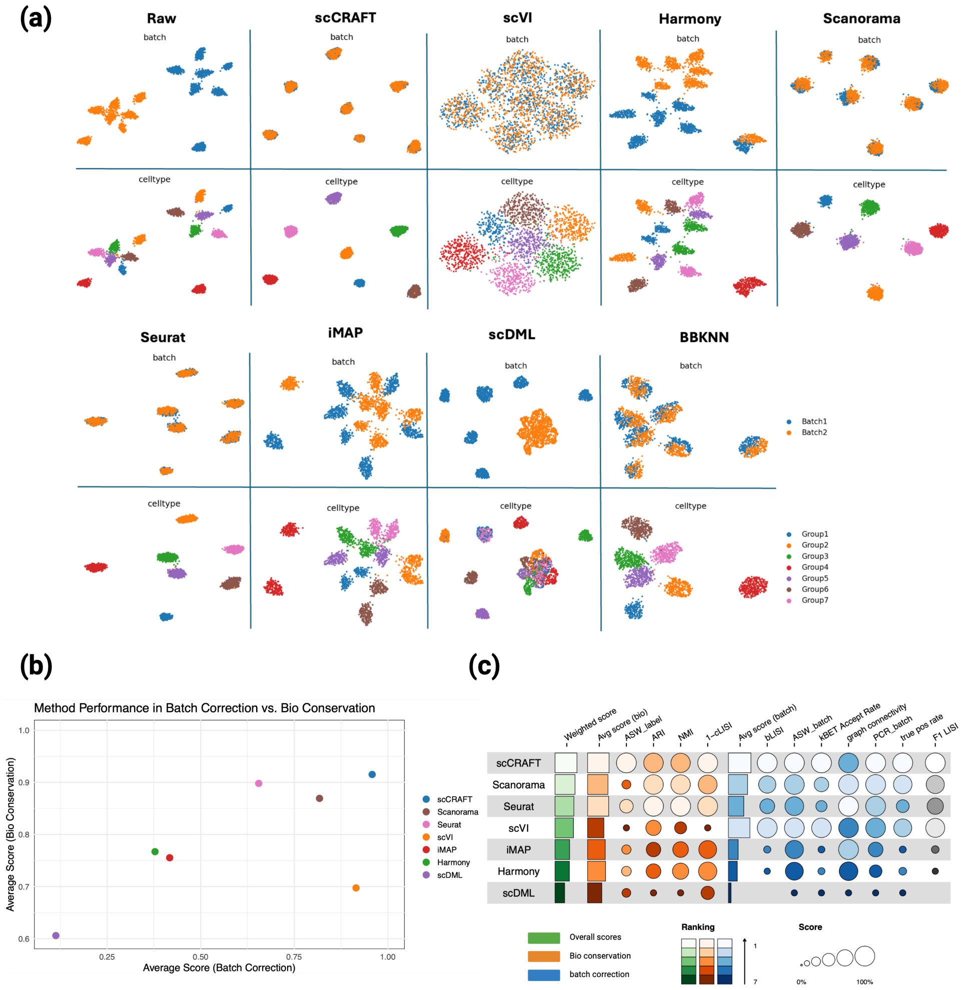

**Fig. S7** Benchmarking Results on simulation task: Rare 0.5. See Suppl Section 6 for details.

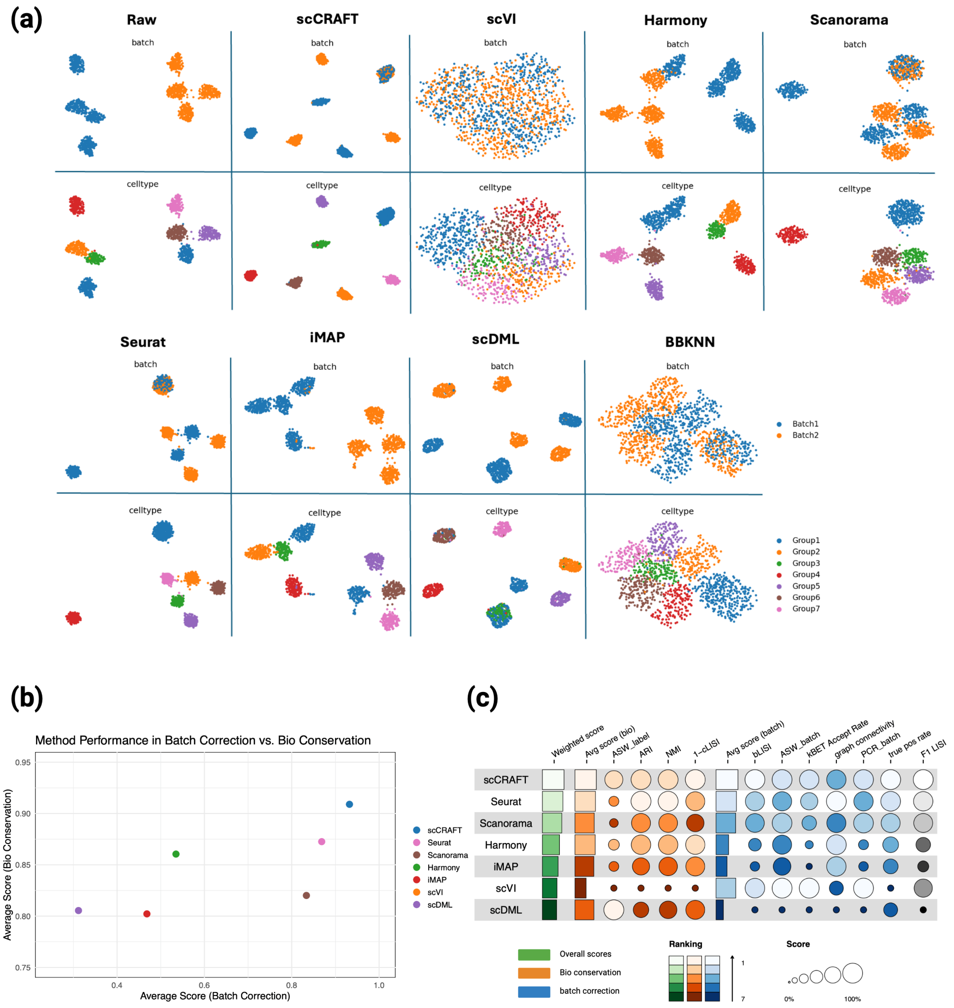

**Fig. S8** Benchmarking Results on simulation task: Common 1. See Suppl Section 6 for details.

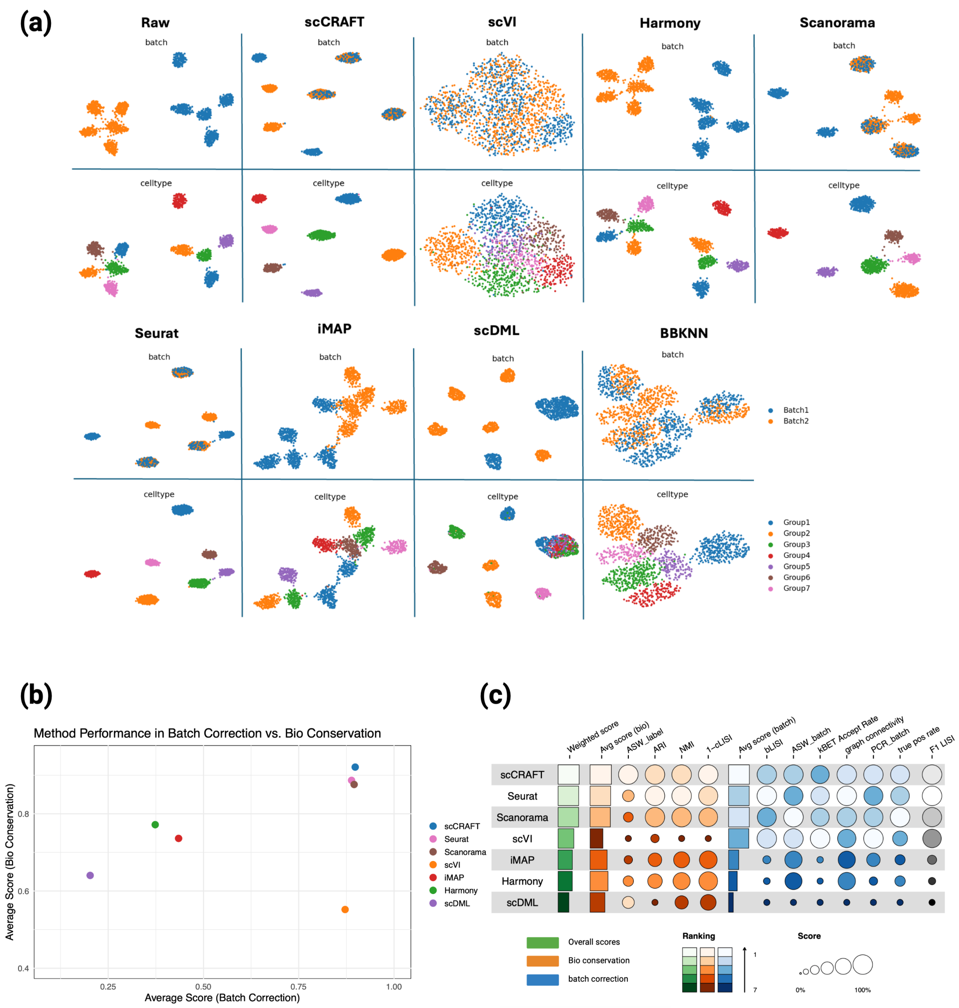

**Fig. S9** Benchmarking Results on simulation task: Common 3. See Suppl Section 6 for details.

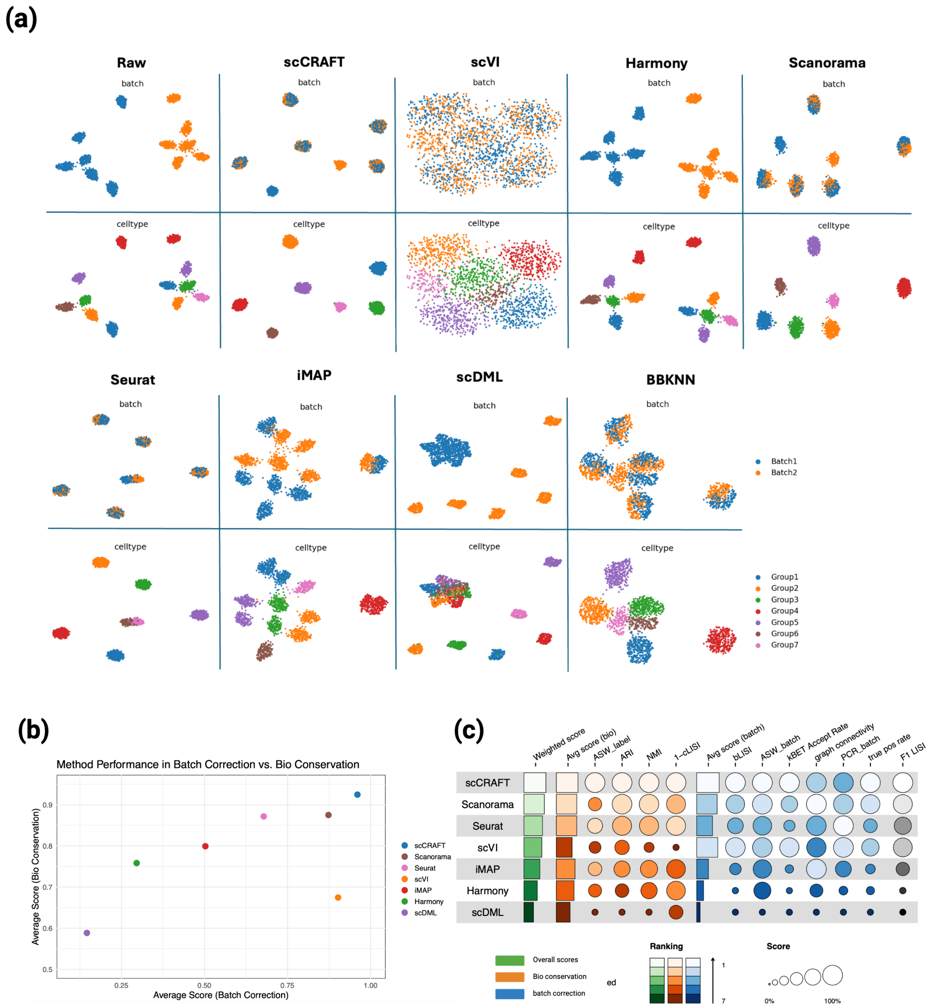

**Fig. S10** Benchmarking Results on simulation task: Common 5. See Suppl Section 6 for details.

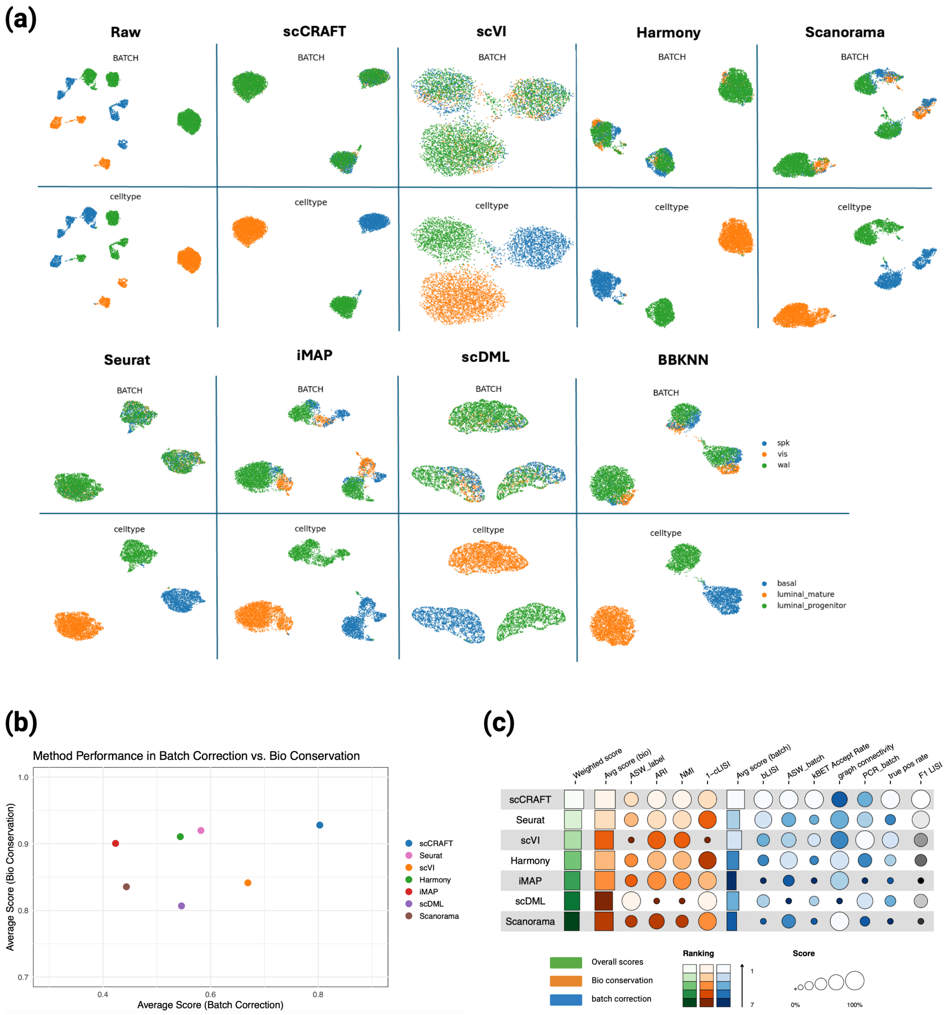

**Fig. S11** Benchmarking Results on real data task: Bct dataset

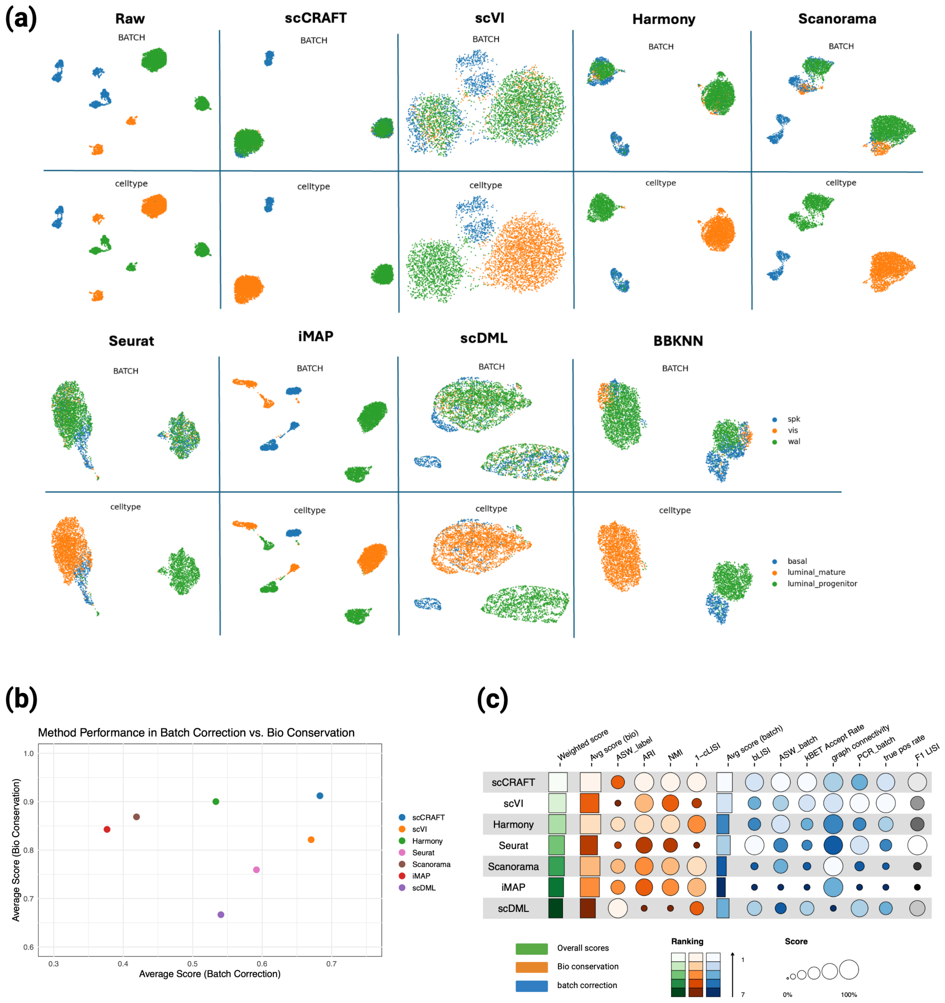

**Fig. S12** Benchmarking Results on real data task: Bct del dataset. See Suppl Section 6 for details.

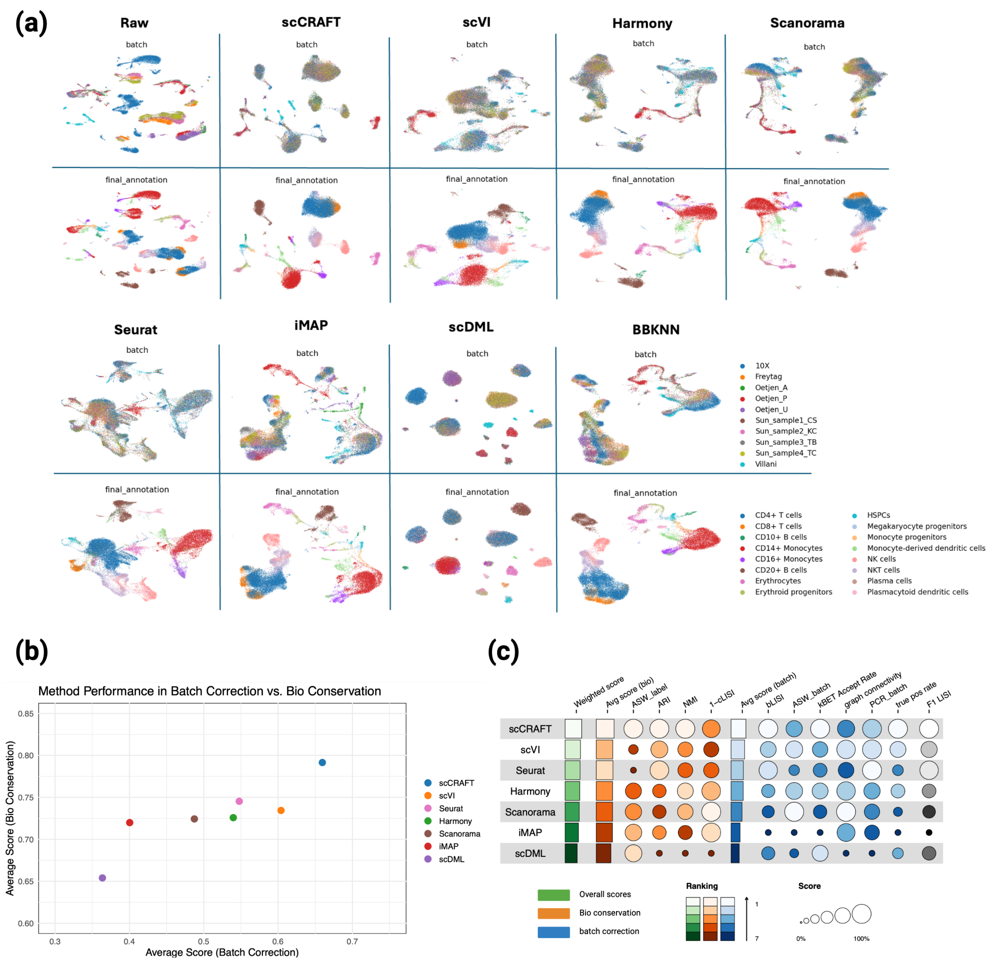

**Fig. S13** Benchmarking Results on real data task: Human Immune

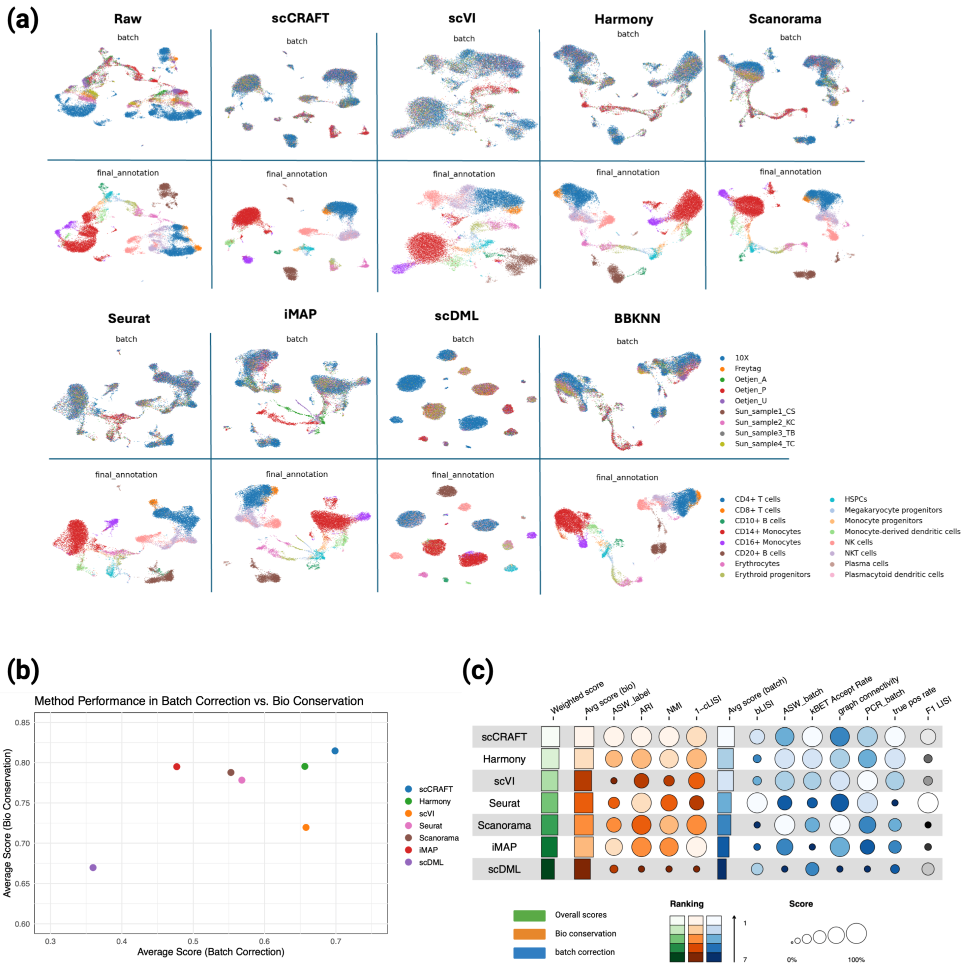

**Fig. S14** Benchmarking Results on real data task: Human Immune Part. See Suppl Section 6 for details.

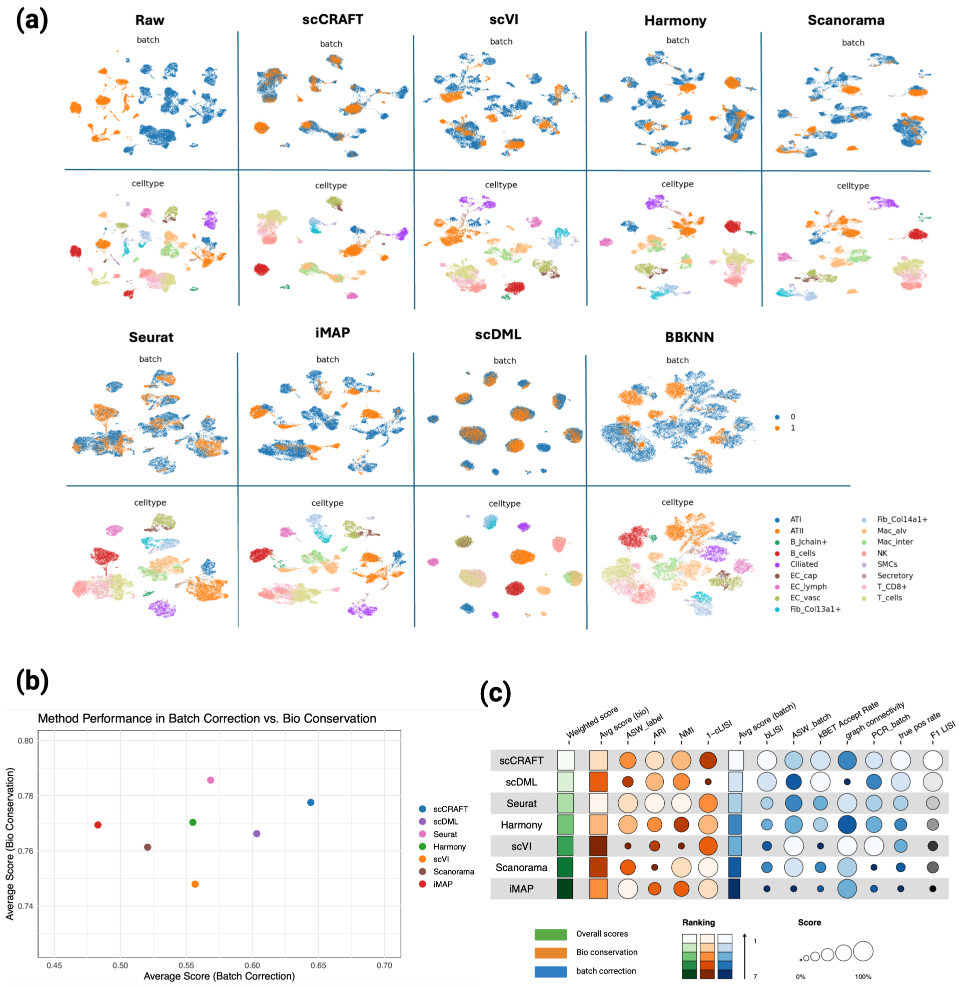

**Fig. S15** Benchmarking Results on real data task: Lung Two Species. See Suppl Section 6 for details.

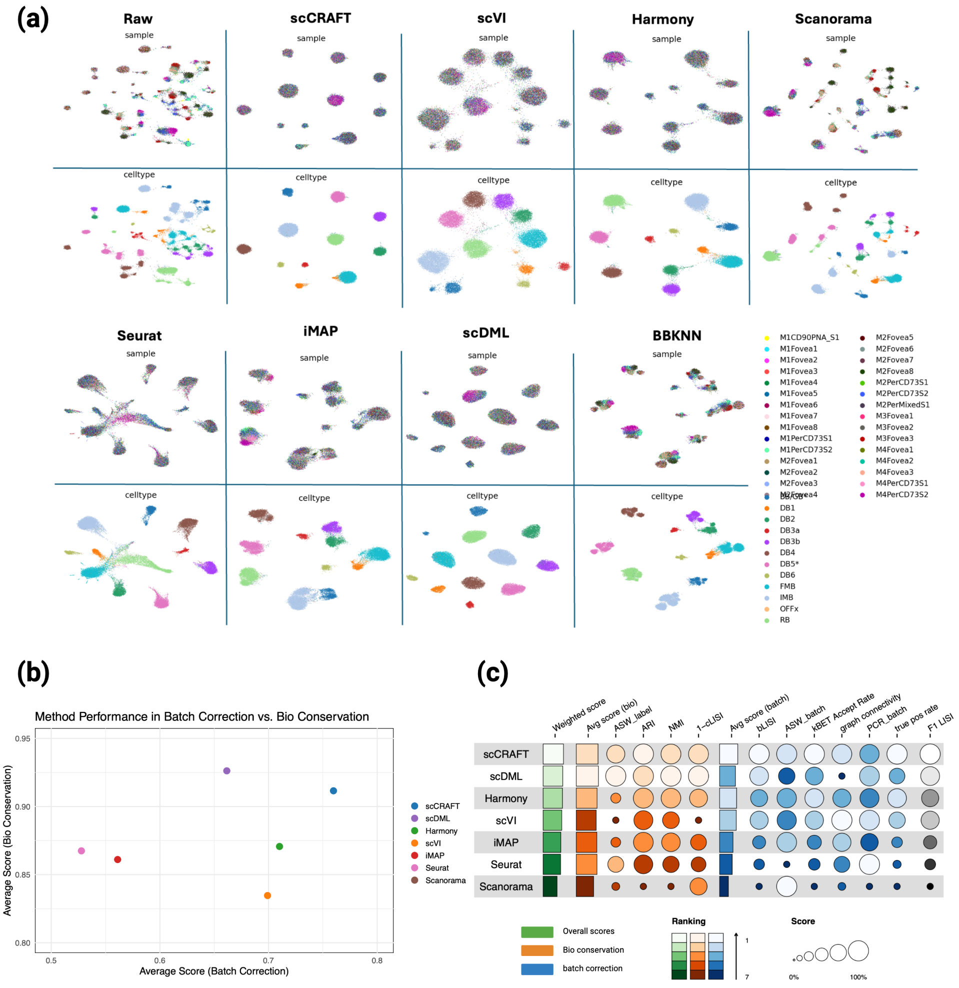

**Fig. S16** Benchmarking Results on real data task: Macaque. See Suppl Section 6 for details.

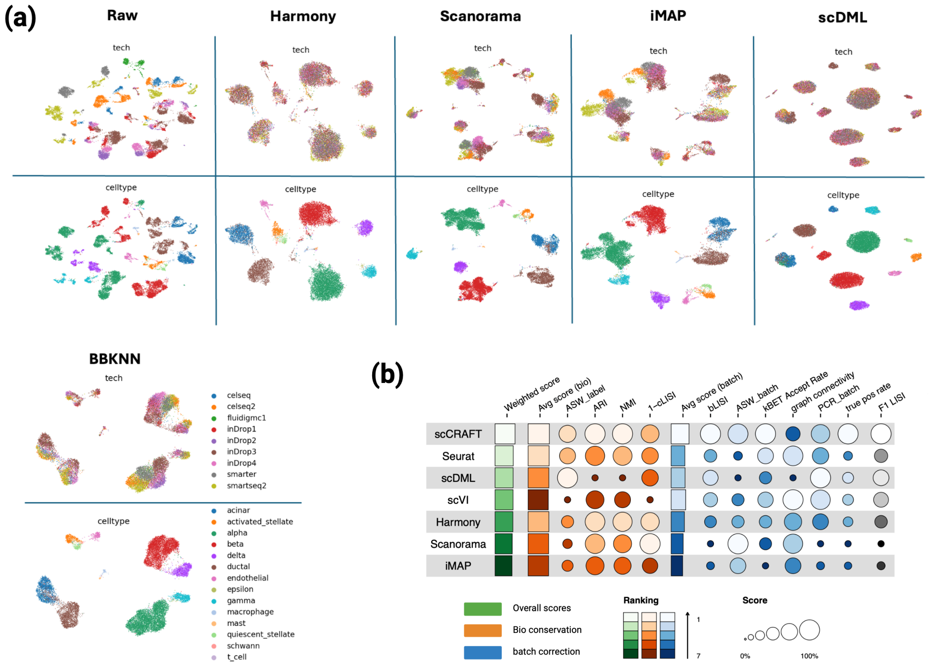

**Fig. S17** Benchmarking Results on real data task: Pancreas. See Suppl Section 6 for details.

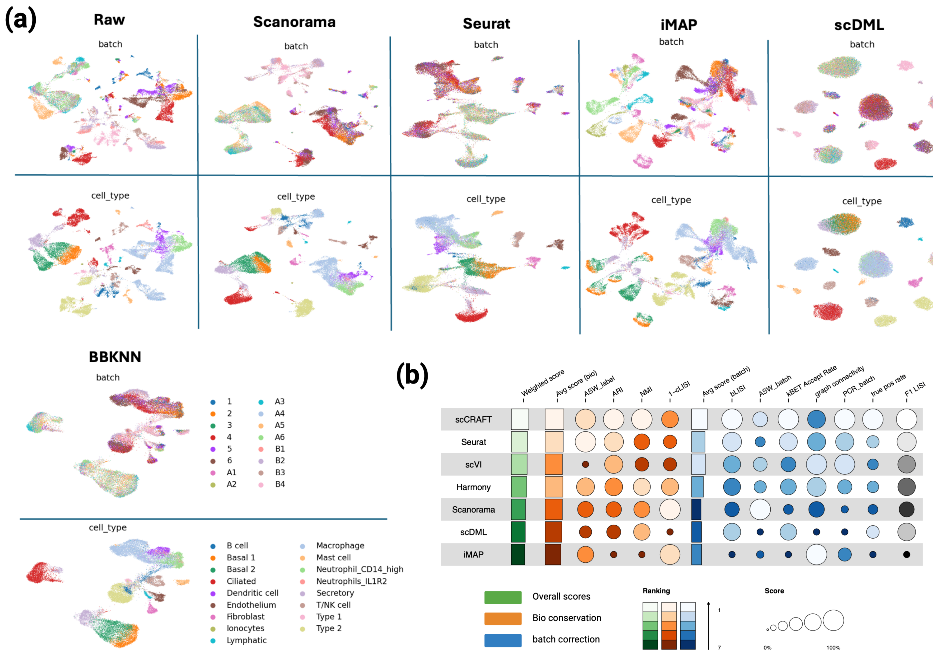

**Fig. S18** Benchmarking Results on real data task: Lung atlas. See Suppl Section 6 for details.

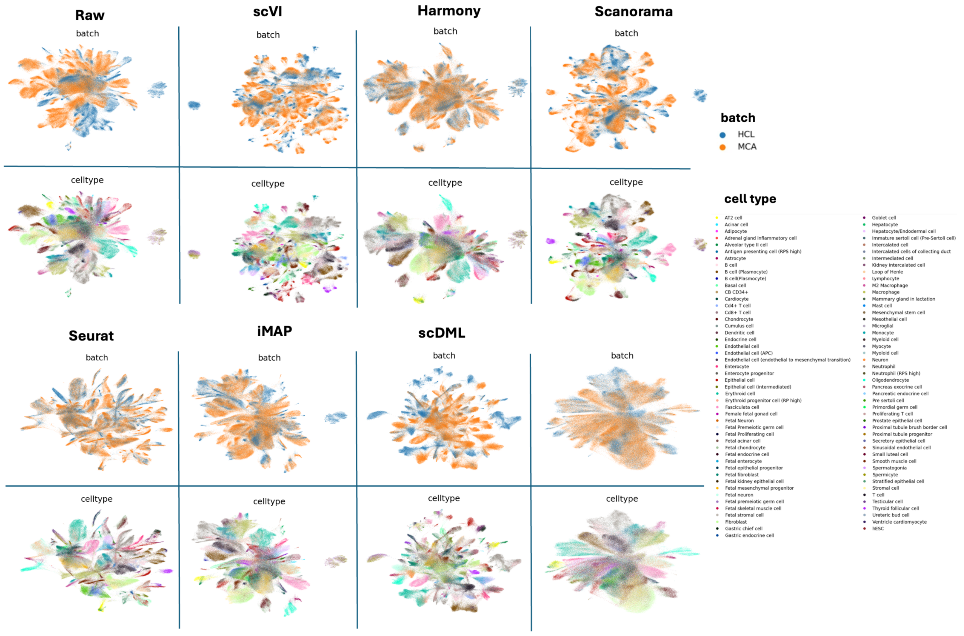

**Fig. S19** Benchmarking Results on real data task: HCL and MCA. See Suppl Section 6 for details.

#### 6.2 Parameter tuning and ablation study

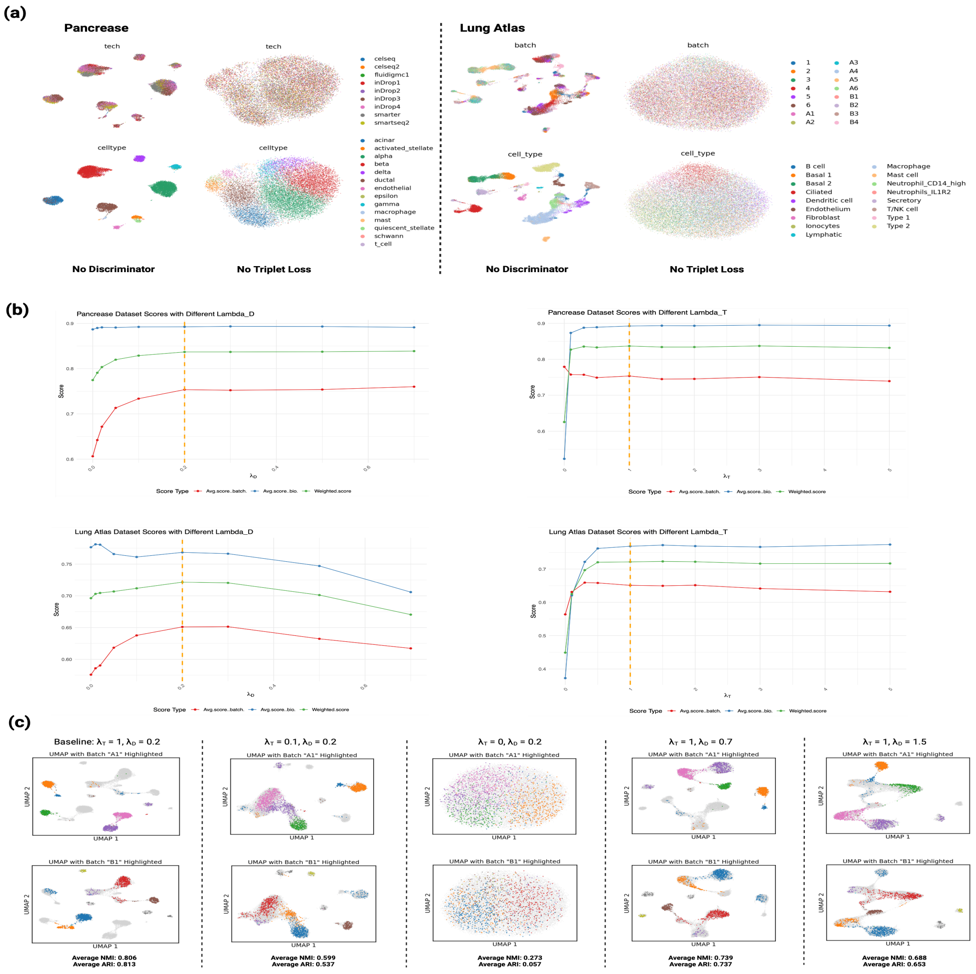

**Fig. S20** Training parameters tuning and ablation study

**(a)** UMAP visualizations detailing the ablation study results for the pancreas and lung datasets, specifically examining the impact of excluding the discriminator or the triplet loss component.
**(b)** Four line plots illustrates the changes in bio-conservation, batch-correction, and overall score for both pancreas and lung datasets when we adjust the parameters $\lambda_T$ and $\lambda_D$ and keep other coefficients unchanged.
**(c)** illustrates the original low-resolution cluster topology retention within a single batch under various parameter settings. The ARI and NMI metrics, calculated using the low-resolution cluster labels, are averaged across all batches to provide a quantitative measure of topology retention across different parameter setups.

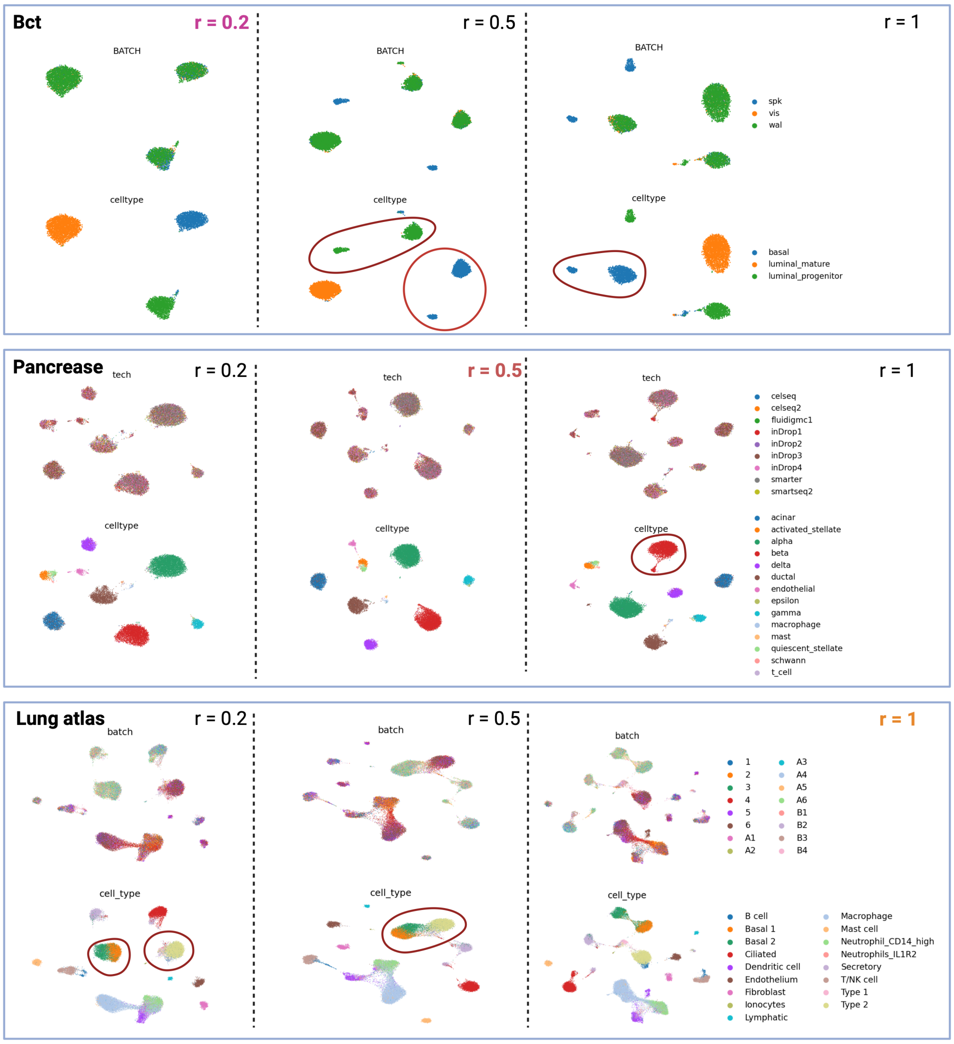

**Fig. S21** Low resolution value tuning.

**(a)** UMAPs for three integrated datasets—bct, pancreas, and lung atlas—under varying low-resolution settings (0.2, 0.5, 1). Each dataset is integrated with different low resolution values, with lighter colors denoting the exact resolutions selected for benchmarking analysis. Red circles within the figure highlight specific conditions under which same cell types were inadvertently divided into separate clusters or different cell types were positioned closely together, indicating potential areas of concern in the integration process
